# Inhibitory control of frontal metastability sets the temporal signature of cognition

**DOI:** 10.1101/2020.08.20.259192

**Authors:** Vincent Fontanier, Matthieu Sarazin, Frederic M. Stoll, Bruno Delord, Emmanuel Procyk

## Abstract

Cortical neural dynamics organizes over multiple anatomical and temporal scales. The mechanistic origin of the temporal organization and its contribution to cognition remain unknown. Here we demonstrate the cause of this organization by studying a specific temporal signature (autocorrelogram time constant and latency) of neural activity. In monkey frontal areas, recorded during flexible cognitive decisions, temporal signatures display highly specific area-dependent ranges, as well as anatomical and cell-type distributions. Moreover, temporal signatures are functionally adapted to behaviorally relevant timescales. Fine-grained biophysical network models, constrained to account for temporal signatures, reveal that after-hyperpolarization potassium and inhibitory GABA-B conductances critically determine areas’ specificity. They mechanistically account for temporal signatures by organizing activity into metastable states, with inhibition controlling state stability and transitions. As predicted by models, state durations non-linearly scale with temporal signatures in monkey, matching behavioral timescales. Thus, local inhibitory-controlled metastability constitutes the dynamical core specifying the temporal organization of cognitive functions in frontal areas.

## Introduction

Large scale cortical networks are anatomically organized in hierarchies of inter-connected areas, following a core-periphery structure (Markov et al., 2013). Within this large scale organization, the dynamical intrinsic properties of cortical areas seem to also form a hierarchy in the temporal domain (Chaudhuri et al., 2014; Murray et al., 2014). The temporal hierarchy arises from increasing timescales of spiking activity from posterior sensory areas to more integrative areas including notably the lateral prefrontal and midcingulate cortex. Intrinsic areal spiking timescales are defined from single unit activity autocorrelation (Murray et al., 2014). Long spiking timescales potentially allow integration over longer durations, which seems crucial in the context of higher cognitive functions, learning and reward-based decision-making (Bernacchia et al., 2011). Recent studies uncovered links between single unit working memory and decision-related activity and spiking timescales in the lateral prefrontal cortex (Cavanagh et al., 2018; Wasmuht et al., 2018). However, the mechanisms that causally determine the timescale of cortical neuron firings and their role in the functional specificity of areas remain to be described.

To address this question, we recorded in the midcingulate cortex (MCC) and lateral prefrontal cortex (LPFC), because these two frontal areas both display particularly long spiking timescales and are functionally implicated in cognitive processes operating over extended timescales. These interconnected regions collaborate in monitoring performance and in integrating the history of outcomes for flexible decisions (Kennerley et al., 2006; Khamassi et al., 2015; Kolling et al., 2018; Medalla and Barbas, 2009; Rothe et al., 2011; Seo and Lee, 2007; Womelsdorf et al., 2014a). Recent anatomical and physiological investigations revealed that the cingulate region has relatively higher levels of synaptic inhibition on pyramidal neurons than LPFC, with higher frequency and longer duration of inhibitory synaptic currents (Medalla et al., 2017), suggesting that excitatory and inhibitory cell types differentially contribute to the specific dynamics of distinct frontal areas. Moreover, MCC also seems to have a longer spiking timescale than the LPFC (Cavanagh et al., 2018; Murray et al., 2014).

In this context, we sought to understand the relationship between temporal features of spiking activity, local neural network dynamics and the computations implemented by frontal neural networks. We focused on whether and how different temporal features play distinct roles in different frontal areas. To this aim, we addressed the following questions: what are the exact differences in the temporal organization of spiking in the LPFC and MCC? How do they relate to the distinct roles of excitation and inhibition? Do they reflect cognitive operations, and can they be adjusted to current task demands? Can they be accounted for by local biophysical circuit specificities? If so, do distinct collective network neurodynamics emerge from such areal biophysical characteristics and what are their functional implications?

We examined the contribution of single unit temporal signatures to dynamical differences between LPFC and MCC in monkeys. After clustering units based on spike shape (putative fast spiking and regular spiking units) we computed spike autocorrelograms and their temporal signatures (time constant and latency). We discovered that LPFC and MCC differed not only in average time constant, but also specifically in the autocorrelogram latency of their regular spiking units.

Regular and fast spiking MCC neurons showed different temporal signatures. Remarkably, through these signatures, neurons contributed to encoding information at different timescales, i.e. information relevant between trials or across multiple trials. Exploring constrained biophysical recurrent network models, we identified the ionic after-hyperpolarization potassium (AHP) and inhibitory GABA-B receptor conductances as critical determinants mechanistically accounting for the difference in spiking temporal signatures between LPFC and MCC. The models predicted how differences in temporal signature amounts to the ability of networks to undergo metastable states with different properties. Indeed, we found, in monkey data, long-lasting states in primate MCC activity but not in the LPFC.

Critically, we show that by controlling states stability and transitions, local inhibition – rather than synaptic excitation (Chaudhuri et al., 2015) – is the major factor setting temporal signatures. Moreover, inhibitory-mediated temporal signatures did not require specific disinhibition between molecularly identified subnetworks of interneurons but naturally emerged from inhibitory weight variability (Wang, 2020).

## Results

We analyzed population spiking timescales for units recorded in MCC and LPFC (140 and 159 units, respectively), using the autocorrelogram of spike counts (see Online Methods), and observed population autocorrelograms similar to those obtained with other datasets (Cavanagh et al., 2018; Murray et al., 2014; Wasmuht et al., 2018) (**Fig. 1a**). At the population level, the characteristic timescale of spiking fluctuation over time, TAU (the time constant from the exponential fit), was longer for MCC than for LPFC (MCC= 519±168 ms, LPFC= 195±17 ms). In addition, MCC single units exhibited longer individual TAUs than LPFC units (medians, MCC=553 ms, LPFC=293 ms; Two-sided Wilcoxon signed rank test on log(TAU), W=15192, p<10^−8^), as in previous datasets (Fig. 1c in Cavanagh et al. (Cavanagh et al., 2018)). Aside from being characterized by a slow decay (long TAU), the MCC population autocorrelation displayed a distinctive feature: a positive slope at the shortest time lags equivalent to a latency in the autocorrelogram, that can be observed in previous publications (see Figure 1c in Murray et al. (Murray et al., 2014), Figure 1d in Cavanagh et al. (Cavanagh et al., 2018)). However, the method we employ above (derived from Murray et al.) cannot resolve the fine dynamics of neuronal activity at short time lags. To improve upon this approach, we instead developed a method based on the autocorrelogram of individual units from all spike times, that provides high temporal precision in parameter estimation (see Online Methods).

**Figure 1.**
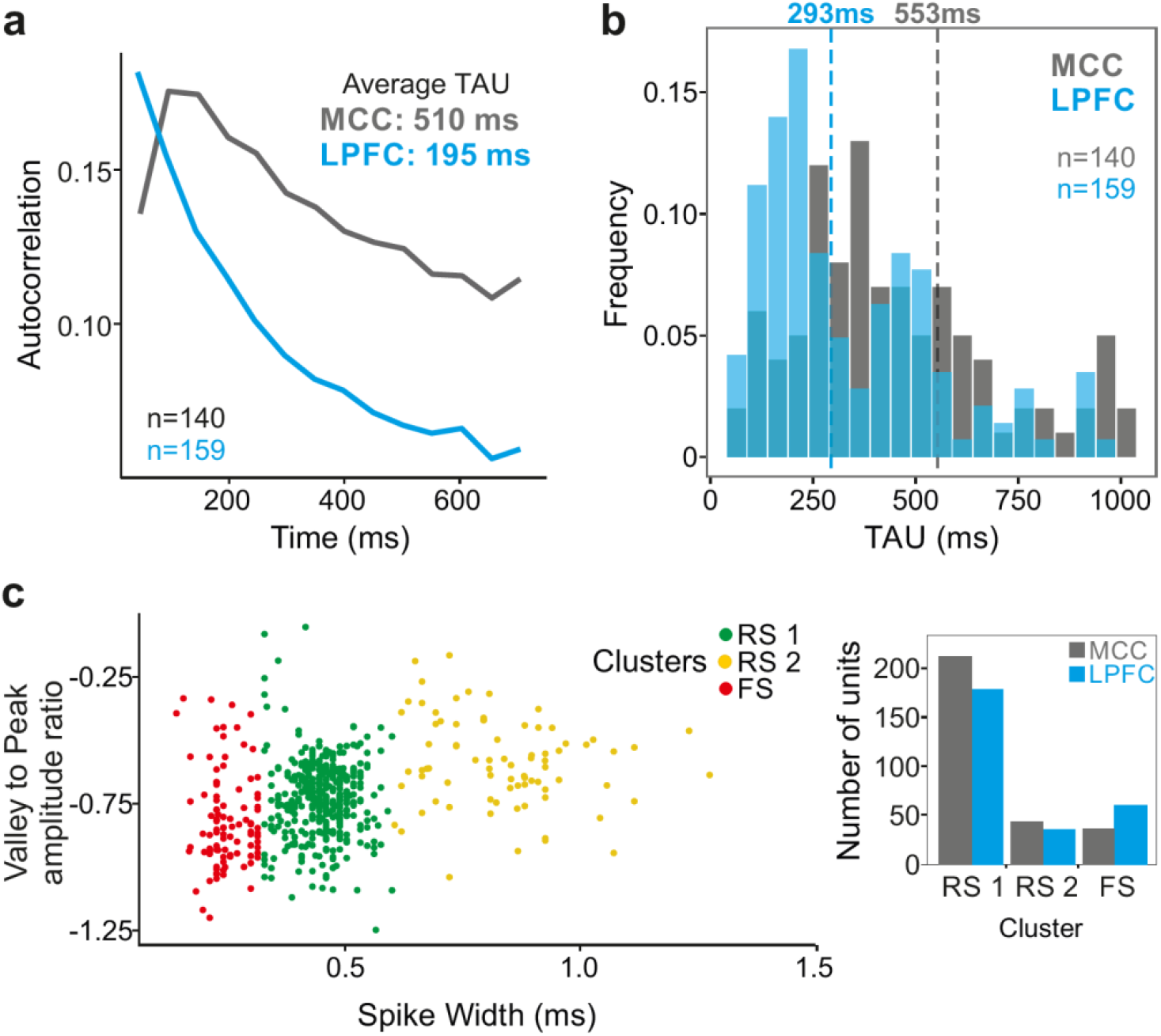
Midcingulate cortex (MCC) and lateral prefrontal cortex (LPFC) spike count autocorrelograms. (**a**) Population exponential fit: autocorrelograms were computed for each unit and the fit was performed on all the units of each area (as in Murray et al. 2014). (**b**) Single unit fits were used to capture individual spiking timescales and produce the distribution of TAU values for each region. Dotted lines represent the median of TAU. (**c**) Clustering of spike shape. We extracted spike width and valley to peak ratio (V2P) from each unit average waveform. A hierarchical clustering led to 3 groups of units (colored groups RS1, RS2, FS). In the paper, units with narrow spike width were termed as fast spiking (FS), whereas units with broader waveform were marked as regular spiking (RS: RS1 + RS2). The histogram indicates the number of MCC and LPFC units belonging to each of the 3 clusters.

One basic assumption to explain local dynamical properties is that interactions between cell types (e.g. pyramidal cells and interneurons) might induce specific dynamics in different areas (Medalla et al., 2017; Wang, 2020; Womelsdorf et al., 2014b). To separate putative cell populations in extracellular recordings we clustered them using single unit waveform characteristics (Nowak et al., 2003). Clustering discriminated 3 populations, with short, large and very large spikes (**Fig. 1c**). The results below were obtained using 2 clusters (small, and large + very large), as detailed analyses showed no clear difference between large and very large spike populations (see **supplementary fig. S1**). We classified units as fast spiking (FS, short spikes; n_MCC_=37, n_LPFC_=61 units) or regular spiking (RS, long spikes; n_MCC_=257, n_LPFC_=215 units) which, in previous studies, were associated to putative interneurons and pyramidal cells respectively.

### MCC temporal signatures differ for regular spiking units

From spike autocorrelograms we extracted multiple metrics: the peak latency (LAT) and time constant (TAU) (see Online Methods). Together, TAU and LAT constituted the temporal signature of single neurons spiking dynamic. The success rate of fitting an exponential on spike autocorrelograms was 91.9% and largely outperformed the alternative method (see Online Methods). **Fig. 2a** shows comparative examples. Note that in the pool of neurons where TAU was successfully extracted using both methods (see method for criteria), we found the two measures (Murray methods vs. spike autocorrelograms) of TAU were correlated (Spearman correlation: rho(282) = 0.46,p<10^−15^). Importantly, TAU was not correlated with firing rate across units (**supplementary fig. S2a**).

**Figure 2.**
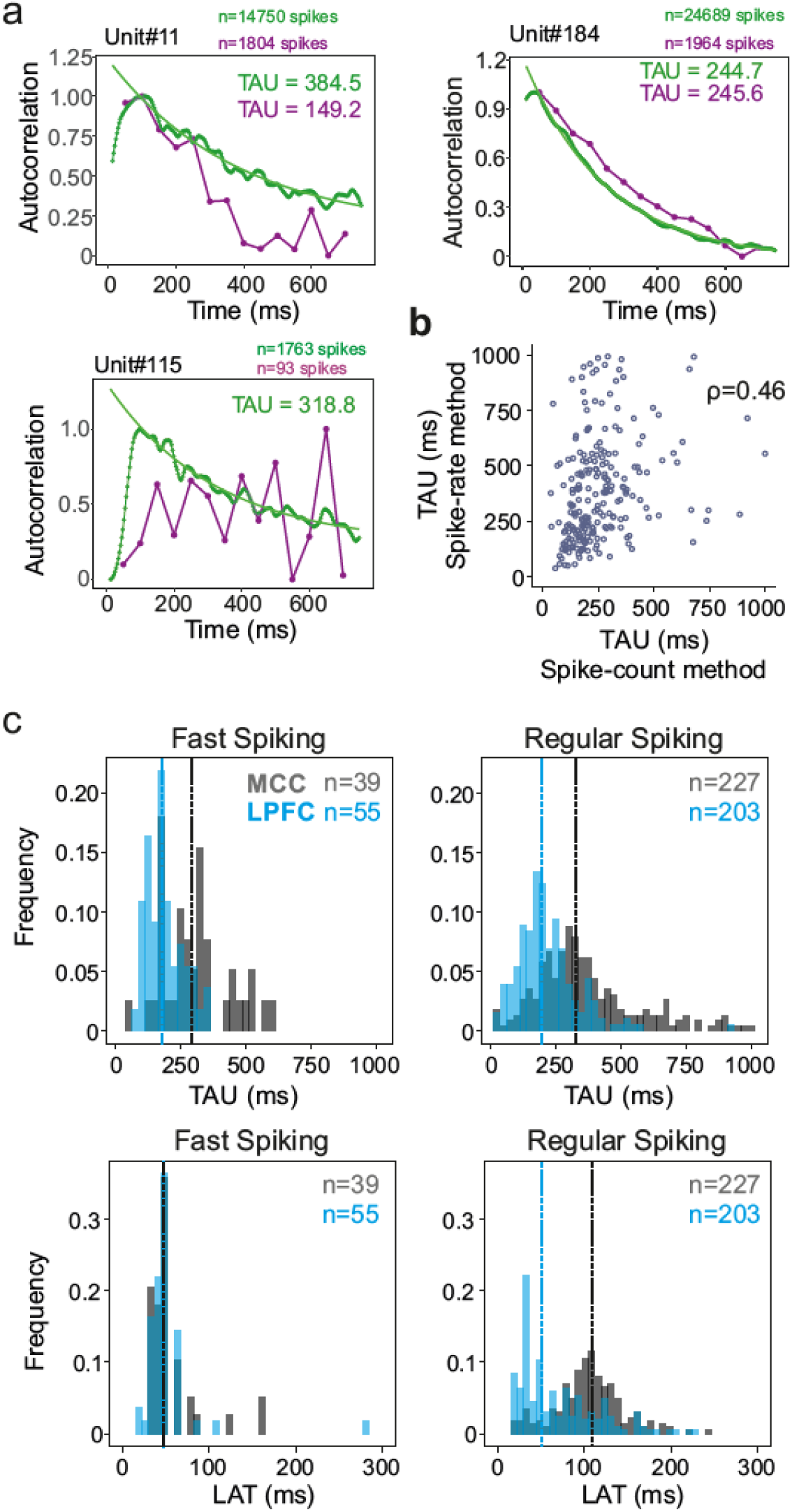
Spike autocorrelogram and temporal signatures in MCC and LPFC. (**a**) 3 single examples of spike count (purple) versus normalized spike autocorrelograms (green) contrasting the outcome of the 2 methods. The measured time constant (TAU) is indicated for both when possible. Numbers of spikes used for each method is also indicated. (**b**) TAU values extracted from each methods are significantly correlated (spearman rho(282) = 0.46,p<10-15). (**c**) Distributions of TAUs (upper histograms) and peak latencies (LAT - lower histogram) for FS (left) and RS (right) units. ‘n’ indicates the number of units. TAU values were longer in MCC than in LPFC for both FS and RS (linear model fit on BLOM transformed TAU for normality, TAU = Region * Unit type, Region: t=−4.68, p<10-6, Unit type: ns, interaction: ns). Peak latencies significantly differed between MCC and LPFC for RS but not for FS units (medians: MCC FS= 48.5 ms, RS= 102.0 ms, LPFC FS= 48.5 ms, RS= 51.8 ms; linear model fit on BLOM transformed Latency for normality, Latency = Region * Unit type, interaction: t-value=−3.57, p< 10-3).

TAU was higher on average in MCC than in LPFC for both regular and fast spiking cells (medians ± sd: MCC FS= 284.7±132 ms, RS= 319.5±199 ms, LPFC FS= 175.1±67 ms, RS= 191.6±116 ms; linear model fit on Blom transformed TAU for normality, TAU = Area * Unit type, Area : F(1,520)=18.36, p<10^−4^, Unit type: F(1,520)=2.72, p=0.12, interaction: F(1,520)=0.19, p=0.79) (**Fig. 2c**).

In addition, LAT became a precise measure obtained for most autocorrelograms. Importantly, it differed significantly between MCC and LPFC for RS but not for FS units, with MCC RS units having particularly long latencies (median ± sd: MCC FS = 48.5±30 ms, RS = 108.7±64 ms, LPFC FS = 48.5±35 ms, RS= 51.9±46 ms; linear model fit on Blom transformed LAT for normality, LAT = Area * Unit type, interaction: F(1,520) = 11.81, p<0.005) (**Fig. 2c**).

TAU and LAT both reflect temporal dynamics, but those measures were significantly correlated only in LPFC RS units (Spearman correlations with Bonferroni correction, only significant in LPFC RS: rho(203) = 0.29, p<10^−3^)). The absence of correlation suggested TAU and LAT likely reflect different properties of cortical dynamics. Moreover, the data suggested that and the different temporal signatures of RS units could reflect differences in the physiology and/or local circuitry determining the intrinsic dynamical properties of MCC and LPFC.

### MCC temporal signatures are modulated by current behavioral state

A wide range of temporal signatures might reflect a basic feature of distributed neural processing (Bernacchia et al., 2011). But do different temporal signatures play distinct roles in terms of neural processing in different areas? And, are these signatures implicated differentially, depending on task demands? As single units were recorded while monkeys performed a decision-making task (described in Stoll et al., 2016; **Fig. 3a**), we extracted each unit’s temporal signature separately for periods in which monkeys were either engaged in the cognitive task or were pausing from performing the task. TAU extracted during engage and pause periods were significantly correlated across neural populations (Pearson correlation: r(267)=0.24, p<10^−4^), indicating that TAU reflects stable temporal properties across conditions. The MCC RS population exhibited a significant modulation of TAU, expressing longer TAU during engage periods compared to pause periods, suggesting that engagement in cognitive performance was accompanied by a lengthening of temporal dynamics for RS neurons in MCC (**Fig. 3b** **left**)(Wilcoxon signed-ranks test (Median=1) with Bonferroni correction, only significant for MCC RS: Median=1.08, V=4265, p<10^−7^). We observed no significant variation of LAT with task demands.

**Figure 3.**
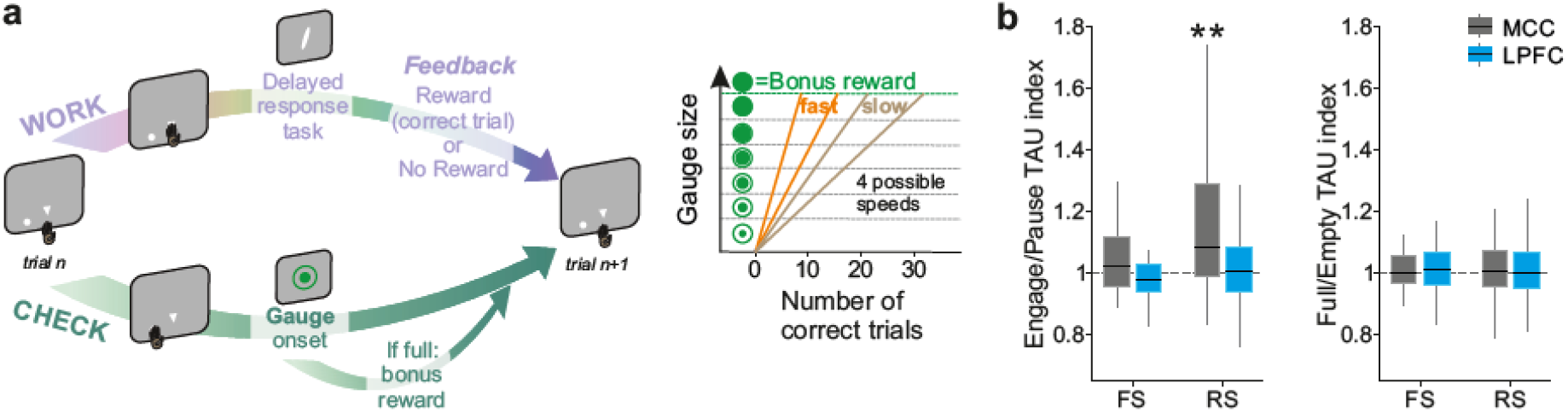
Behavioral engagement in task and spiking timescale changes. (**a**) Schematic representation of the task. At the start of each trial, animals can either initiate a delayed response task (WORK option) which can lead to 1 reward delivery, or use the CHECK option to check the current size of the gauge (or collect the bonus reward). Each reward in the task contributes to increase the gauge size and bring the bonus availability closer. The graph (right) schematized the speed of increase of the gauge size which varies between blocks (fast or slow blocks). (**b**) Boxplots of indices for each unit type and region calculated to estimate potential changes in TAU between Engage and Pause (left), and between empty and full gauge (right). TAUs increased in Engage vs. Pause only for MCC RS units.

### Temporal signatures are linked to cognitive processing

Contrary to MCC, LPFC temporal signatures were not modulated by engagement in the task. Multiple cognitive models propose a functional dissociation between MCC and LPFC and indeed empirical data reveal their relative contribution to feedback processing, shifting, and decision making (Khamassi et al., 2015; Kolling et al., 2018; Stoll et al., 2016). One important question is thus whether temporal signatures observed for a given area and/or cell type contribute to selected aspects of cognitive processing. For example, temporal signatures might be adjusted to the current functional context and time scale required to perform a task. In our experiment monkeys gained rewards by performing trials correctly in a categorization task while each success (reward) also brought them closer to obtaining a bonus reward (**Fig. 3a**, right panel, see Online Methods for task description). By touching a specific lever at trial start, animals could either enter a categorization trial or check the status of a visual gauge indicating the proximity of the bonus reward availability. The number of rewards (i.e. correct categorization trials) needed to get the bonus, and thus the speed of the gauge increase, varied across blocks (i.e. either fast or slow). Previous analyses revealed that feedback influenced the likelihood of checking in the following trial (Stoll et al., 2016). Thus, feedback can be considered as information used on a short timescale (within the inter trial period). The animals also built an estimation of the gauge size that was updated upon checking in order to regulate the frequency of checks during blocks, allowing animals to seek and collect the bonus in a cost-efficient manner (Stoll et al., 2016). Gauge size can thus be considered as information used and carried over long timescales.

We first hypothesized that blocks of different speeds and/or gauge encoding could engage neurons and modulate their spiking timescale. This was not the case. TAU values were not significantly modulated depending on the state of the gauge (less vs. more than half full, **fig. 3b** **right**), nor related to different speeds (Wilcoxon signed-ranks test (Median=1) with Bonferroni correction, for gauge state and gauge speed, all p>0.6).

Conversely, we assessed whether temporal signatures observed for certain cell types contributed to code specific aspects of the task. We used mixed effect models on groups of single units to test the contribution of population activity to encoding task relevant information: feedback in categorization trials (i.e. reward vs. no-reward), and gauge size. The rationale was that feedback information was relevant within the intertrial period, whereas Gauge information was relevant across trials between two successive checks. Previous analyses had revealed that both MCC and LPFC units encode such information, although MCC units showed greater contributions(Stoll et al., 2016). We classed both FS and RS units as either short *or* long TAU units using a median split. A time-resolved generalized mixed linear models (*glmm*) revealed notable dissociations between these populations. During the intertrial period, the population of MCC RS units with short TAU was mostly involved in encoding feedback information, which was relevant only for the current trial (**Fig. 4a**). By contrast, RS units with long TAU were mostly involved in encoding gauge information, which contributed to regulate decisions across trials (Stoll et al., 2016) (**Fig. 4a**, lower right). Long and short TAU RS populations in LPFC contributed mostly to encode feedback during the intertrial period (Fig. 4a, right).

**Figure 4.**
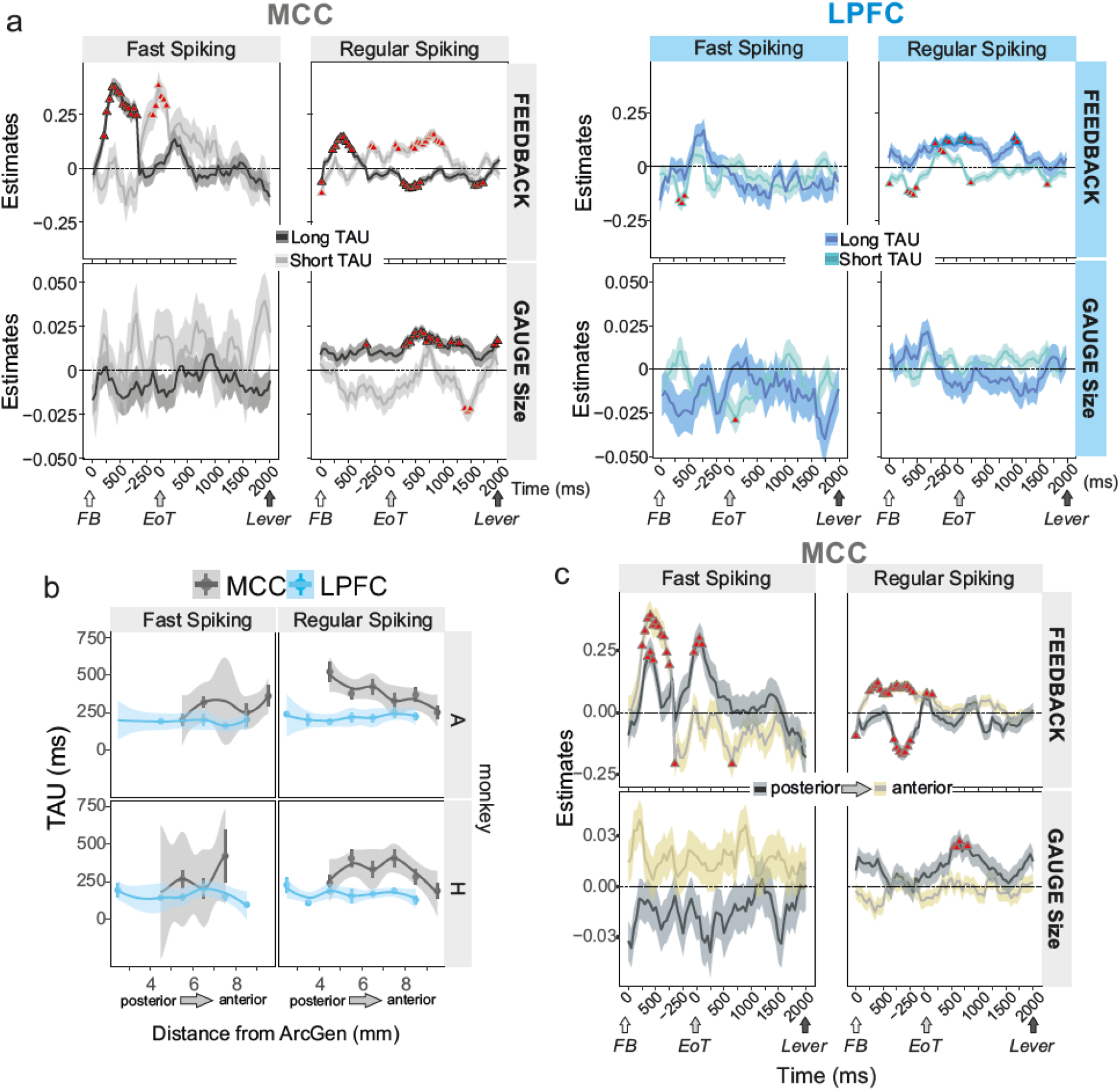
Encoding of feedback and gauge size for different unit types and spiking timescales and rostro-caudal distribution. (**a**) Estimates (β-coefficients) obtained from the MCC (grey) and LPFC (blue) unit populations obtained from time-resolved *glmm* for Feedback (reward vs no reward; top graphs) and Gauge size (bottom) (see ‘Group analyses using *glmm*’ in Methods). Estimates are obtained at successive time points covering the entire inter-trial period between feedback onset and the lever onset in the following trial. Significant effects are indicated by a red triangle (p<0.05 corrected), shadings indicate standard deviations. Positive values depict a population activity bias towards negative feedback (top) and positive slope of linear coding for gauge size (bottom). Data is presented for FS and RS units (left and right respectively for each panel) and have been performed on subpopulation with short or long TAU values (determined by a median split). Short and long TAU populations are represented by light and dark color intensity. Note in particular the dissociation for RS MCC units with short and long TAU respectively coding for feedback and gauge size. (**b**) Averaged TAU values along the postero-anterior axis in the MCC and LPFC, for both monkeys. (**c**). Estimates reflecting coding strength of Feedback and Gauge size for MCC unit populations separated by their rostro-caudal location.

Interestingly FS units in the MCC were mostly engaged in the first second after feedback onset, with a strong bias toward encoding negative feedback (Fig. 4a, upper left, positive estimates). Effects were more transient and involved short TAU units in the LPFC (**Fig. 4a**).

### Spiking timescales are anatomically organized in MCC

Spiking timescales measured in MCC and LPFC covered several orders of magnitudes (10-1000 ms; **Fig. 2c**). Because single unit recordings spanned large regions, such wide range could reflect anatomical organization of segregated populations with distinct homogeneous intrinsic properties. Such organization has been observed in MCC with human fMRI (Meder et al., 2017). We indeed found that average TAU values in MCC were higher in more posterior parts, in particular for RS units (ANOVA on Blom transformed TAU: MCC, monkey A: F(5,112)=2.8, p=0.041, monkey H: F(5,54)=3.09, p=0.033, LPFC, monkey A: F(6,110)=0.34, p=1, monkey H: F(6,64)=2.49, p=0.066; linear regression on Blom transformed TAU: MCC, monkey A: t(1,112)=8.99, p=0.0067, monkey H: t(1,54)=2.22, p=0.28, LPFC, monkey A: t(1,110)=1.09, p=0.60, monkey H: t(1,64)=0.25, p=1; all p-values are FDR corrected for n=2 comparison per monkey) (**Fig 4b**). This suggests an antero-posterior gradient of spiking timescales. No such effect was observed in LPFC. Similar analyses for LAT revealed no consistent inhomogeneity within MCC or LPFC (**Fig. S2b**).

The consequence of such an organization, knowing the respective functional involvement of units with long and short TAU (**Fig 4a**), should be an antero-posterior functional gradient. We tested this by separating MCC cells in posterior versus anterior subgroups and tested their contribution to feedback and gauge encoding (**Fig. 4b**). Indeed, posterior RS units’ activity contributed to positive encoding of gauge size, preceded in time by encoding of positive feedback (negative estimates) (**Fig. 4c** **lower and upper right**), while anterior RS units showed primarily a contribution to feedback encoding (**Fig. 4c** **upper right**). Finally, anterior FS units were primarily (in time and in strength) contributing to encoding negative feedback. This remarkable contribution of FS to feedback encoding is studied and discussed further below.

In summary, MCC regular spiking units with relatively short or long TAU contributed to the encoding of task elements relevant over short and long terms, respectively. The spiking timescales seemed to be organized along the rostro caudal axis in MCC. This suggests a correspondence between cell type, temporal signatures and their functional involvement in processing specific aspects of cognitive information in different functional subdivisions of cortical regions. The crucial questions thus remain of the mechanistic origin of temporal signatures and of how they relate to cognitive functions.

### Biophysical determinants of temporal signatures in frontal network models

To uncover the source and consequences of distinct temporal spiking signatures in the LPFC and MCC, we designed a fine-grained model of local recurrent frontal networks. This model is unique in combining 1) highly-detailed biophysical constraints on multiple ionic channels, synaptic receptors and architectural frontal specificities, and 2) the cardinal realistic features of mammals cortical neurodynamics including the excitation/inhibition balance, high-conductance state of neuronal activity and asynchronous irregular regime characterizing the awake state (Brunel, 2000; Destexhe et al., 2003; Hennequin et al., 2017). Our specific goal was to evaluate whether biophysical circuit specificities could mechanistically account for differences in LPFC and MCC temporal signatures. We also assessed whether these specificities induce distinct collective network neurodynamics and functional implications, possibly explaining the empirical relationships between temporal signatures, cell type, and information processing.

We first explored, using Hodgkin-Huxley cellular models (see *Online Methods*), whether specific frontal temporal signatures may arise from ionic or synaptic properties of individual neurons. Extensive explorations of these models identified, among many ionic and synaptic conductances tested, the maximal cationic non-specific (g_CAN_) and potassium after-hyperpolarization (g_AHP_) conductances as the sole couple affecting both LAT and TAU. However, their regulation could not fully reproduce the monkey data set (see Supplementary **Fig. S4** and **S5**). Thus, we then assessed whether collective dynamics at the level of recurrent networks models could better account for frontal temporal signatures (**Fig. 5a**, see *Online Methods*). One-dimensional explorations of the large parameter space failed to identify single biophysical determinants accounting, alone, for differences between LPFC and MCC (RS and FS) temporal signatures (Supplementary **Fig. S6 and Table S1**). However, these explorations targeted four parameters of interest regulating either LAT or TAU confirming those already revealed in cellular models (g_CAN_ and g_AHP_) and uncovering, in addition, NMDA and GABA-B maximal conductance (g_NMDA_ and g_GABA-B_) whose slow time constants strongly affected network dynamics.

**Figure 5.**
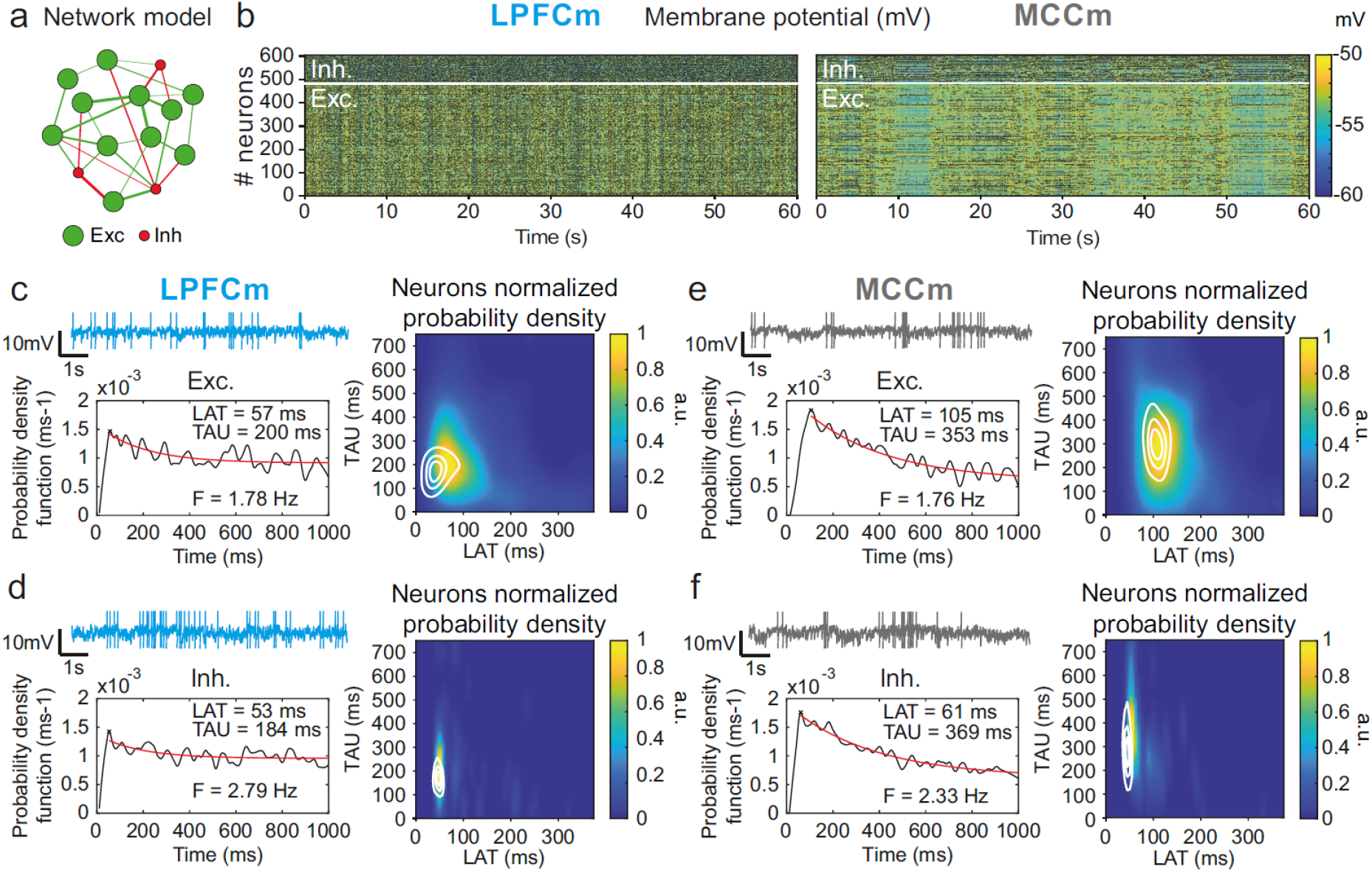
Temporal signature of LPFCm and MCCm recurrent network biophysical models. (**a**) Scheme of the frontal recurrent networks modelled, with 80% excitatory (green) and 20% inhibitory (red) neurons and sparsity of synaptic connections. (**b**) Membrane potential in the 484 excitatory (lower part) and 121 inhibitory (upper part) neurons of example network models with parameter set to approximate LPFC dynamics (g_CAN_=0.025mS.cm^−2^, g_AHP_=0.022mS.cm^−2^, g_GABA-B_=0.0035mS.cm^−2^; see text and legend of Fig. 6b for the choice of LPFC and MCC standard g_AHP_ and g_GABA-B_ maximal conductances) and MCC dynamics (g_CAN_=0.025mS.cm^−2^, g_AHP_=0.087mS.cm^−2^, g_GABA-B_=0.0143mS.cm^−2^). (**c**) (upper left) Membrane potential of an example excitatory neuron in the LPFC model (LPFCm). Scaling bars 1s and 10mV (spikes truncated). (lower left) Autocorrelogram of this LPFCm example excitatory neuron (black) and its exponential fit (red, see *Online Methods*). (right) Bivariate probability density distribution of autocorrelogram parameters in LPFCm excitatory neurons. Contour lines at 50, 75 and 90% of the maximum of the bivariate probability density distribution in LPFC monkey RS units. (**d**) Same as **(c)** for LPFCm inhibitory neurons, with contour lines from the bivariate probability density distribution in LPFC monkey FS units. **(e,f)** Same as **(c,d)**, for the MCCm and MCC.

Two-dimensional explorations using these key parameters (**Fig. 5** and supplementary **Fig. S7**) identified a single specific setup which demonstrated network dynamics that reproduced the shift from the LPFC-like temporal signature to that resembling the MCC with striking precision. An increase of both g_AHP_ and g_GABA-B_, in the presence of g_CAN_, drove the model from an LPFC-like temporal signature (LPFCm model) (**Fig. 5c & d**; map and contours: bivariate probability density model and monkeys’ distributions, respectively) towards that of the MCC (MCCm model, **Fig. 5e-f**). Specifically, g_AHP_ increased LAT and decreased TAU in excitatory (likely equivalent to RS) neurons (**Fig. 6a** left) and had no effect in inhibitory (likely FS) neurons (**Fig. 6a** right). Besides, g_GABA-B_ decreased LAT in both excitatory and inhibitory neurons (**Fig. 6a** top) and increased TAU in an intermediate range (**Fig. 6a** bottom). A bivariate probability density-based similarity measure (see *Online Methods*) revealed that monkey temporal signatures were robustly reproduced by the model in two large contiguous regions in the (g_AHP_, g_GABA-B_) plane, with both conductances increased in the MCC (**Fig. 6b**).

**Figure 6.**
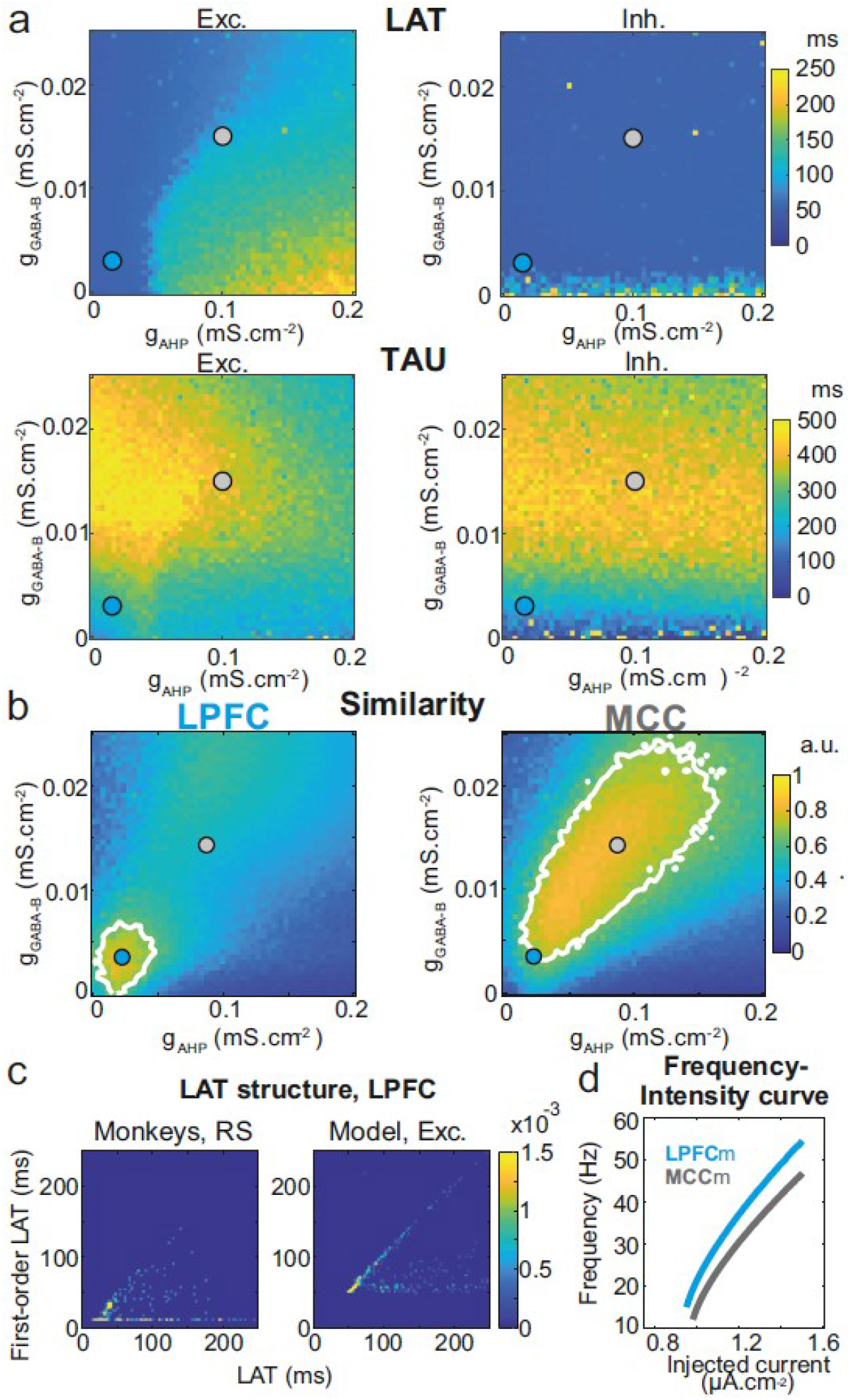
Similarity to monkey LPFC and MCC temporal signatures critically depends on AHP and GABA_B_ conductance in the network model. (**a**) Mean population LAT (top) and TAU (bottom) in Exc (left) and Inh (right) neurons, as a function of AHP and GABA-B maximal conductances. Blue and grey disks indicate the (g_AHP_, g_GABA-B_) parameter values of the LPFCm and MCCm models, respectively. (**b**) Similarity of the temporal signature between the network model and monkey data in the LPFC (left) and MCC (right), as a function of AHP and GABA-B maximal conductances (see Online Methods). In (a) and (b), the value for each (g_AHP_, g_GABA-B_) is averaged over 5 simulations. Contour line at 80% of maximum similarity. LPFCm and MCCm (g_AHP_, g_GABA-B_) parameter values calculated as coordinates of the contour delimited area’s weighted average. (**c**) Bivariate probability density distribution of the autocorrelogram LAT and first- order latency (the latency of the ISI distribution) in RS units in monkey LPFC (left) and excitatory neurons in the example LPFCm model (right). The model accounts for two distinct neuronal subsets in RS neurons, where LAT is determined by first-order latency solely (due to gAHP-mediated refractoriness; diagonal band), or in conjunction with other factors (g_GABA-B_ slow dynamics-mediated burstiness and recurrent synaptic weight variability; horizontal band). (d) Single excitatory neuron frequency/intensity relationship in the LPFCm (blue) and MCCm (grey) models in response to a constant injected current.

Several lines of evidence further indicated the model’s relevance. First, the model properly accounted for the larger LAT variability in monkey RS vs FS units (**Fig. 5**). Moreover, it reproduced the complex relations between LAT and first-order latency (ISI distribution latency) remarkably well, in all populations (**Fig. 7c** and **Supplementary Fig. 7**). Furthermore, both the firing frequency and input-output gain were lower in MCCm excitatory neurons (**Fig. 6d**), because of its higher g_AHP_ (Naudé et al., 2012), as found experimentally(Medalla et al., 2017).

**Figure 7.**
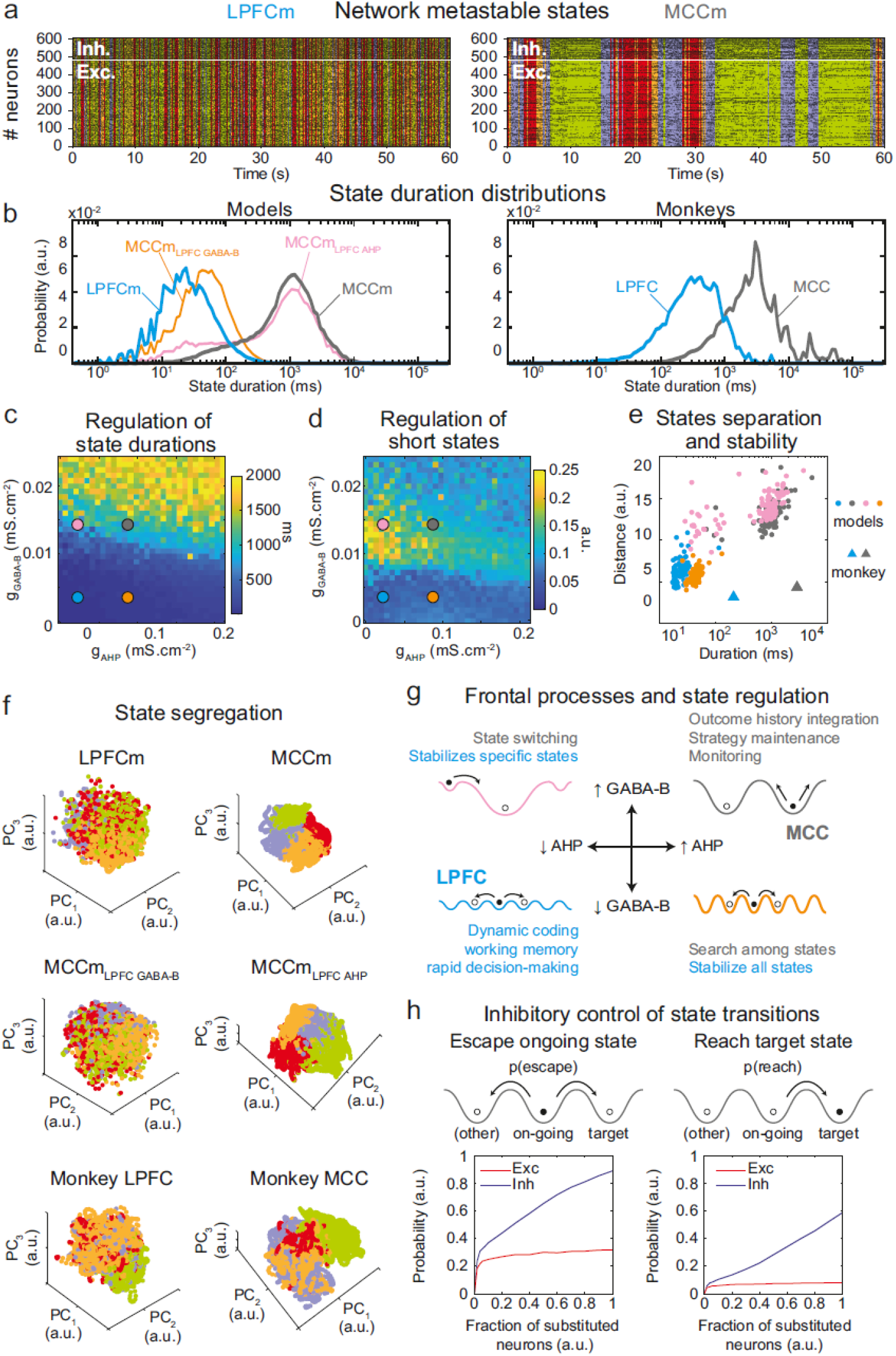
Properties of metastable states in the LPFC and MCC. (**a**) LPFCm and MCCm models spiking raster plots (black dots), with Hidden Markov model states (HMM, colored bands) (**b**) State duration distributions: probability distributions of being in states of given durations in LPFCm (blue), MCCm (grey), MCCm with LPFCm g_AHP_ (MCCm_LPFC AHP_ pink) and MCCm with LPFCm g_GABA-B_ (MCCm_LPFC GABA-B_, orange) models (left) and monkey LPFC (blue) and MCC (grey) areas (right). Each model was simulated 100 times and analyzed via HMM, while monkey data was analyzed via HMM with 100 different initiation parameter states. Periods above 300s were excluded. (**c, d**) Regulation of state duration and short states: median state duration (c) and Kolmogorov-Smirnov one-sample test statistic or maximal distance of state duration probability distributions to log-normality, as a measure of the over-representation of short states (d), as a function of g_AHP_ and g_GABA-B_ maximal conductances. Colored disks indicate parameter values of LPFCm, MCCm, MCCm_LPFC AHP_ and MCCm_LPFC_ GABA-B models, respectively. Each point is the average of 5 simulations. (**e**) Separation between states: average distances between HMM states (averaged pairwise distance between neural centered standardized frequency centroids (temporal averages) of HMM states), as a function of median state durations. Distances calculated over 100 simulations in models and once for monkey LPFC and MCC data. (**f**) State segregation: projection of neural activity on the principal components of the PCA space of example model simulations and of monkey data. State colors as in (a). (**g**) Frontal processes and state regulation: schematic attractor landscapes in the LPFC and MCC. Horizontal and vertical arrows indicate possible regulations of AHP and GABA_B_ conductance levels respectively by intrinsic/synaptic plastic processes or neuromodulation in the LPFC and MCC. Likely functional processes operating in these landscapes are indicated in blue for the LPFC and grey for the MCC. (**h**) Inhibitory control of state transitions: probability to escape an ongoing state (left) and to reach a target state (right), when the ongoing state is perturbed by substituting a given proportion of its excitatory (vs inhibitory) neurons’ activity by that of the same neurons in the (perturbing) target state (see Online Methods). Average (full line), +/− s.e.m. (shaded areas, almost imperceptible).

### Metastable states underlie LPFC and MCC temporal signatures

The asynchronous irregular (presumably chaotic) dynamics of network models was highly structured in time (**Fig. 5b**). Hidden Markov models (HMM) revealed that it organized through collective transitions between so-called metastable (quasi-stationary) states in the models LPFCm and MCCm (**Fig. 7a**), as found in frontal areas (Abeles et al., 1995; Seidemann et al., 1996; Xydas et al., 2011). Moreover, while LPFCm states maximally lasted a few hundred milliseconds (**Fig. 7b**, **left, blue**), MCCm states persisted up to several seconds (**Fig. 7b**, **grey**). This suggested that such a difference in metastability may also parallel the difference of temporal signature in monkey LPFC and MCC areas. Applying HMM to experimental data revealed that, as predicted by the model, neural activity was organized as metastable states at slower timescales in the MCC (vs the LPFC, **Fig. 7b**, **right**). State durations were globally shorter in models (compared to monkeys), as they contained neither temporal task structure nor learning (see discussion) and were not optimized to fit data.

Long states essentially required high g_GABA-B_ in the MCCm, as they disappeared when g_GABA-B_ was lowered to its LPFCm value (MCCm_LPFC GABA-B_ model, **Fig. 7b** left, orange curve). In contrast, they only marginally depended on g_AHP_. MCCm and an MCCm model with the g_AHP_ derived from that of LPFCm (MCCm_LPFC AHP_) showed state duration distributions that were essentially similar, although there was a small increase in the probability of short states at lower g_AHP_ (pink vs gray curves). In the (g_AHP_, g_GABA-B_) space, g_GABA-B_ systematically proved to be essential in increasing the duration of states, with a border region that clearly separated short states (<0.1s) from longer states (>1s) (**Fig. 7c**) At this intermediate border, lower g_AHP_ increased the probability of short states (grey vs pink dots; distributions were even bimodal at lowest g_AHP_ values, not shown), as witnessed by departure from log-normality (**Fig. 7c**). As such, the temporal structure of states in the LPFCm was dominated by short and unimodal state duration distributions (**Fig. 7c** and **7d**, blue dots), as in monkeys (**Fig. 7b**, right) and previous studies(Abeles et al., 1995; Seidemann et al., 1996). In the MCCm, by contrast, the distribution displayed large durations and a slight departure from log-normality (**Fig. 7c** and **7d**, grey dots), resulting in a majority of long states (>1s) coexisting with short states, as found in data (**Fig. 7b**).

State duration, i.e. stability, scaled with spatial separation in the neural space of activity (**Fig. 7e**, see Online Methods). Indeed, the shorter states of network models with lower g_GABA-B_ (LPFCm and MCCm_LPFC GABAB_, blue and orange dots) were less distant, compared to those of networks models with higher g_GABA-B_ (MCCm and MCCm_LPFC AHP_, grey and pink dots). While states were largely intermingled in the LPFCm and MCCm_LPFC GABAB_ (**Fig. 7f**, upper & middle left), they clearly segregated in the MCC and MCCm_LPFC AHP_ (**Fig. 7f**, upper & middle right). As predicted by the model, segregation between states was indeed higher in the monkey MCC (**Fig. 7e**, large grey triangle, and **Fig. 7f**, lower right), compared to the LPFC (**Fig. 7e**, large blue triangle, and **Fig. 7f**, lower left). This suggests that the higher stability of states in monkey MCC arose from a larger segregation of representations in the space of neural activity.

Altogether, these results suggested that itinerancy between metastable states constitutes a core neurodynamical principle underlying the diversity of computational processes and functions operated in primate frontal areas (**Fig. 7g**, see Discussion). From this perspective, the conditions governing transitions between states is critical. We thus evaluated how perturbations of selective neuronal populations would escape ongoing states and reach specified target states (**Fig. 7h**). In the MCCm, we substituted the membrane potentials and synaptic opening probabilities of a fraction of excitatory (vs inhibitory) neurons of the ongoing HMM state by those of a target state. This could mimic the effect of internal chaotic fluctuations or external inputs aimed at reaching that target state. Surprisingly, escaping the ongoing state or reaching the target state remained quite unlikely when substituting excitatory neurons, whatever the fraction (**Fig. 7h**, left). By contrast, both probabilities of escaping and reaching scaled with the fraction of substituted inhibitory neurons, with high maximal probabilities (mean: 0.89 and 0.59 for escaping and reaching, respectively – **Fig. 7h**, right panel). Interestingly, the probability of escaping a state could attained 0.24 even with as few as 2% of substituted inhibitory neurons, indicating the significant impact of single inhibitory neurons on state itineracy.

Thus, inhibition is a major factor controlling targeted transitions between metastable states in the MCC network model and is also crucial in determining their stability. Excitation had no such role. This result is remarkable, especially considering that MCC FS neurons encoded negative outcomes immediately after feedback onset that triggered behavioral adaptive responses (**Fig. 4**). This could reflect the involvement of MCC FS neurons in inducing state changes on feedback associated to behavioral flexibility.

## Discussion

We showed LPFC and MCC displayed long population spiking timescales (TAU), with larger values in MCC (TAU~500 vs 200 ms), consistent with previous observations (Chaudhuri et al., 2015; Murray et al., 2014). In fact, LPFC and MCC express distinctive and complex temporal organizations of their activity, which cannot be solely captured by the population spiking timescale. The spiking timescale has been used as a measure characterizing intrinsic areal properties and an inter-area temporal hierarchy. However, the spiking timescale of single units varied over two orders of magnitude within each area (Cavanagh et al., 2018; Murray et al., 2014; Wasmuht et al., 2018). The latency of autocorrelogram also demonstrate informative variability, which suggest important underlying functional richness. Our study demonstrates that the temporal signature (TAU and LAT) of single units, measured through spike autocorrelogram metrics and cell type segregation, can highlight specific local ionic and synaptic mechanisms. Differences in temporal signatures, for instance between LAT of FS and RS in MCC, and within regions, provide important information on the functional properties of the underlying neural network.

Unravelling the multidimensional nature of LPFC and MCC temporal signatures at the level of individual neurons enabled us to constrain refined biophysical recurrent network models and reveal the local biophysical determinants mechanistically accounting for their specific temporal organization. Moreover, we showed that these determinants control neurodynamical features that constitute core computational foundations for the executive cognitive processes operated by these frontal areas.

### Functional spatio-temporal organization of temporal signatures in frontal areas

The correlation between temporal signatures and behavior suggests how such biophysical properties could contribute to functional specificities. Spiking timescales distributions have been related to persistent activity, choice value and reward history in the LPFC and MCC (Bernacchia et al., 2011; Cavanagh et al., 2018; Meder et al., 2017; Wasmuht et al., 2018). Here, the spiking timescales of MCC RS units increased on average during periods of engagement in cognitive performance, likely reflecting the global implication of neural processes in task performance at long timescales. MCC units with different temporal signatures differentially contributed to cognitive processes known to engage MCC, namely feedback/outcome processing and outcome history representations (Kennerley et al., 2009; Quilodran et al., 2008; Seo and Lee, 2007). Outcome processing generally enables rapid – trial by trial – adaptation of control and decisions, while outcome history representations contribute to the long-term – across trials – establishment of values guiding strategy adaptation (Behrens et al., 2007; Karlsson et al., 2012).

In our experiment, short spiking timescale units contributed to feedback processing, whereas long spiking timescale units and especially RS units, contributed to encode gauge size, which linearly increase with the accumulation of rewards across trials. In MCC, this temporal dissociation coincided with a spatial organization along the antero-posterior axis: anterior units mainly encoded feedback valence, more strongly and earlier than posterior units, whilst posterior units mostly encoded the long-term information related to gauge size. This antero-posterior gradient strikingly resembles that observed in humans (Meder et al., 2017).

### Local molecular basis of frontal temporal signatures

Through extensive parameter exploration of constrained biophysical frontal network models, we identified 2 conductances that precisely reproduced all monkey temporal signatures. In the model, higher TAU (i.e. MCC vs LPFC, posterior vs anterior MCC) was accounted for by stronger synaptic GABA-B levels, consistent with reported higher GABA-B receptor densities (Zilles and Palomero-Gallagher, 2017), stronger and slower inhibitory currents in the MCC (vs LPFC) (Medalla et al., 2017), and stronger GABA-B receptor densities in the posterior (vs anterior) MCC (Palomero-Gallagher et al., 2009). Excitatory synaptic transmission has been proposed to be a crucial determinant of longer spiking timescales in the temporal cortical hierarchy (Chaudhuri et al., 2015). We found that while stronger excitatory transmission increases TAU (possibly accounting for longer MCC TAUs), it also decreases LAT. LAT, however, was longer in the monkey MCC. This suggests that GABA-B inhibitory – rather than excitatory – transmission is the causal determinant of longer spiking timescales, at least in the LPFC and MCC. Noticeably, long timescales do not require specific disinhibition between molecularly identified subnetworks of interneurons (Wang, 2020) but naturally emerge from inhibitory weights variability (see below). The model also predicts that higher LAT in the MCC originate from increased refractoriness through higher after-hyperpolarization potassium (AHP) conductances in RS units. Higher AHP implies lower input-output gains in MCC RS units, compared to the LPFC (Naudé et al., 2012), as found empirically (Medalla et al., 2017). Finally, reproducing appropriate temporal signatures required the cationic non-specific (CAN) conductance in the areas’ RS units. This was observed in RS of rodent medial frontal areas (Haj-Dahmane and Andrade, 1997; Ratté et al., 2018), where it regulates, together with AHP, cellular bistability and memory, network persistent activity and computational flexibility (Compte, et al., 2003; Papoutsi et al., 2013; Rodriguez et al., 2018; Thuault et al., 2013). Our conclusions do not preclude the contribution of other factors to temporal signatures such as large-scale hierarchical gradients (Chaudhuri et al., 2015), distinct neuromodulations (see below), or inputs with different spectral contents to LPFC and MCC.

### Frontal temporal signatures uncover metastable dynamics

The LPFC and MCC activity, both in models and in monkeys’, was metastable, i.e. organized in sequences of discrete, quasi-stationary states in which activity fluctuates around fixed-point attractors (Abeles et al., 1995; La Camera et al., 2019; Rich and Wallis, 2016; Seidemann et al., 1996). As a general rule, the duration of states increases with the stability of their attractor (i.e. the depth/width of their basin of attraction) and decreases with spiking fluctuations. Fluctuations originate from stochastic inputs or chaotic noise (as in our model), and they trigger state transitions.

States were longer in monkeys, likely because extensive training induced attractors that were more stable, whereas models displayed less stable attractors that simply resulted from just random connectivity without learning. Thus metastability genuinely emerged from synaptic heterogeneity and did not require strong network clustering (La Camera et al., 2019). We showed that high GABA-B levels are crucial to stabilize states because they amplify the heterogeneity of inhibition and widens attractors, as reflected by higher state separation in the MCC. In addition, GABA-B’s long time constant naturally promotes burstiness, i.e. stable discharge episodes. Finally, higher AHP levels, required for higher LAT in MCC RS units, limited the occurrence of the shortest states, limiting frequent transitions between states.

In monkeys and biophysical models, temporal signatures, which correlate with state stability, actually reflect the underlying temporal organization of neurodynamics into metastable states. Interestingly, state durations (up to >10s) were longer than spiking timescales (<0.5s), reconciling the apparent discrepancy between typical spiking timescales in frontal areas (<1s) and the functional timescales at which those areas operate (up to tens of seconds, Bernacchia et al., 2011).

### Functional significance of controlled metastable states in frontal areas

Metastable states can be linked to specific representations in the brain at a variety of levels of abstraction, from stimuli to mental states (Engel et al., 2016; La Camera et al., 2019; Mazzucato et al., 2015, 2019; Rich and Wallis, 2016; Taghia et al., 2018). In general, state transitions contain appreciable randomness, with high transition rates signing internal deliberation, whilst more stable states predicting forthcoming decisions (La Camera et al., 2019). We suggest that controlling itinerancy among metastable states constitutes a core neurodynamical process supporting executive functions in frontal areas, which allows to scan choices and strategies, generate deliberation and solve on-going tasks.

Specifically, in the MCC (**Fig. 7g**, gray landscape) GABA-B-mediated long metastable states underlying long spiking timescales may contribute to the maintenance of ongoing strategies (Durstewitz et al., 2010; Enel et al., 2016; Stoll et al., 2016) and to the integration of outcome history (Kennerley et al., 2006; Meder et al., 2017; Seo and Lee, 2007; Tervo et al., 2014). At shorter timescales, short states might instantiate dynamic coding, flexible computations and rapid decision-making in the LPFC (Fig. 7g, blue landscape) (Rich and Wallis, 2016; Rigotti et al., 2013; Stokes, 2015). Short states may be lengthened in the LPFC when AHP is increased (Fig. 7g, orange landscape), favoring longer timescales and a global stabilization of, for instance, working memory processes (Cavanagh et al., 2018; Durstewitz and Seamans, 2008). Conversely, decreasing GABA-B destabilizes all long states in the MCC model, globally favoring fast transitions (Fig. 7g, orange landscape). This mechanism might contribute to abandon prior beliefs and to rapid search for adapted representations, e.g. in uncertain environments (Karlsson et al., 2012; Quilodran et al., 2008; Stoll et al., 2016). In the LPFC model with increased GABA-B or in the MCC model with decreased AHP, activity destabilizes certain long states, favoring transitions to remaining long states (Fig. 7g, pink landscape). Such a configuration might be relevant for flexible behaviors, directed exploration and switching (Durstewitz et al., 2010; Pasupathy and Miller, 2005; Russo et al., 2020; Stoll et al., 2016). Regulating GABA-B and AHP to dynamically adapt computations and temporal signatures could be achieved through neuromodulatory or fast plastic processes (Froemke, 2015; Satake et al., 2008).

Macroscopic gradients of inhibitions and excitations appear as important determinants of the large scale organization of cortical dynamics (Wang, 2020; Womelsdorf et al., 2014b). Our results indicate a complementary fundamental dual role of local inhibition in regulating state durations and stability on one hand, and setting the timing and direction of state transitions, on the other. Moreover, transitions can be easily triggered using very few inhibitory neurons. Our study suggests that interneurons and inhibition might be causal in error-driven state transitions in the MCC. Such transitions, initiated by FS neurons immediately after feedback onset, would allow escaping currently unsuccessful states, reaching alternatives or exploring new states.

In conclusion, we showed that local ionic and synaptic determinants specify the scale of temporal organization of activity in frontal cortical areas. These determinants might produce the particularly long states observed in monkey MCC dynamics and could explain its contribution to functions operating over extended behavioral periods. More generally, our results suggest that the diversity of spiking timescales observed across the cortical hierarchy reflects the local excitability- and synaptic inhibition-mediated regulation of metastability, which sets the temporal organization of computational processes.

## Materials and methods

### Subjects and materials

This project was conducted with two male rhesus monkeys (Macaca mulatta), monkey A and H. All procedures followed the European Community Council Directive (2010) (*Ministère de l’Agriculture et de la Forêt, Commission nationale de l’expérimentation animale*) and were approved by the local ethical committee (*Comité d’Ethique Lyonnais pour les Neurosciences Expérimentales*, CELYNE, C2EA #42). Electrophysiological data were recorded using an Alpha-Omega multichannel system (AlphaOmega Engineering, Israel).

### Recording sites

Recording chambers (Gray Matter research, MT, USA) were centered on antero-posterior coordinates of +34.4 and +33.6 relative to ear bars (for monkey A and H, respectively)(Stoll et al., 2016). MCC recording sites covered an area extending over 10mm (anterior to posterior), and at depths superior to 4mm from cortical surface (corresponding to the anatomically defined aMCC or functionally defined dACC). Recording sites in LPFC were located between the principalis and arcuate sulcus (areas 6DR, 8B, 8A and 9/46) and at depths inferior to 2mm from cortical surface. Reconstructions of cortical surface, of MRI sections perpendicular to recording grids and of microelectrode tracks were performed using neuronavigation. Locations were confirmed with MRI reconstructions and stereotaxic measurements by keeping track of electrophysiological activity during lowering of electrodes.

### Single unit activity and spike shapes

Electrophysiological activity was recorded using epoxy-coated tungsten electrodes (1–2MOhm at 1 kHz; FHC Inc., USA) independently lowered using Microdrive guidance (AlphaOmega Engineering). Neuronal activity was sampled at 22 kHz resolution. Single units were sorted offline using a specific toolbox (UltraMegaSort2000, Matlab toolbox, Kleinfeld Lab(Hill et al., 2011), University of California, San Diego, USA). Metrics served to verify the completeness and purity of single unit activity. Each single unit activity was selected, recorded and included in analyses on the basis of the quality of isolation only. We obtained 298 MCC units and 272 LPFC units while monkeys performed a checking task(Stoll et al., 2016). A subset of these data has been used in a previous publication(Stoll et al., 2016).

#### Spike shape clustering

Spike shapes can be clustered in different groups that might correspond to different putative cell populations. For each single unit, we computed the average spike shape on which we measured:

1. Pre-valley (V1): the minimum value of the waveform prior to the peak
2. Post-valley (V2): the minimum value of the waveform following the peak
3. Spike width: the time between the occurrence of the peak and V2
4. The ratio of V1 to V2 (V1/V2)
5. The ratio of V2 to the spike peak (V2/PiK)

We clustered average units according to their spike width and V2/PiK. We first computed the spike width *vs.* V2/PiK Euclidean distance matrix (*dist* function in R). Then we performed hierarchical clustering using Ward’s method (*hclust* function in R). The number of retained clusters was determined with the combination of data viewing, dendrogram examination and objective measures of clustering quality (Elbow method, Average silhouette method and Gap statistic method). The partitioning led to 3 clusters, one with narrow spike shapes, one with wide spikes and one with very wide spikes. Narrow and wide spikes were considered FS and RS, respectively. Although clustering revealed 3 clusters, no differences were found between the 2 wide ones, both considered RS neurons (see supplements).

### Spiking timescales

The primary analysis of timescales was based on Murray et al(Murray et al., 2014). Spike counts were measured in 14 successive bins of 50ms from the pre-cue period (700ms) of each trial, when the monkey is in a controlled, attentive state awaiting stimulus onset. We first calculated the cross-trial bin cross-correlations. Each vector of spike counts from the 50ms bin *t* was correlated with vectors of spike counts at subsequent bins (*t*+1, *t*+2, etc) generating an autocorrelation matrix. The positive side of the autocorrelation was used to compute timescales. The autocorrelogram data was then fitted using non-linear least square (*nls* function in R) to a function of the form:

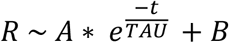

where R is the correlation coefficient and *t* the bin time. TAU, representing the decay of the exponential function and thus the intrinsic timescale, and A, a scaling constant, were obtained from the fit. We computed TAU both at the population level, by using a global fit on all recorded units from a given area (as in Murray et al(Murray et al., 2014)), and at the single unit level.

However, the above method cannot resolve the fine dynamics of neuronal activity at short time lags because it is based on counts pooled across trials and from coarse-grained time bins (50ms). Moreover, the large variability of unit discharge resulted in a high variability of autocorrelograms, which could not be fitted in many cases (47.5% failures), as in other studies (52.1% and 48.4% failures in Wasmuht et al. 2018 and Cavanagh et al. 2018(Cavanagh et al., 2018; Wasmuht et al., 2018), respectively). Finally, tracking the causal determinants of LPFC and MCC temporal signatures in terms of local cellular and/or network dynamics requires a high temporal precision, because they rely on intrinsic and synaptic time constants, which often lie below the coarse time bin of the spike count method. To prevent these shortcomings, we directly computed the autocorrelogram of individual neurons from spike times, allowing for high temporal precision in parameter estimation. For this we leveraged all the data recorded for each neuron to reduce the large noise present at the level of individual neurons.

### Autocorrelogram analysis

To capture the dynamics of neuronal activity, we computed autocorrelograms from individual unit spike timeseries and extracted their latencies (LAT) and time constants (TAU). The same method was applied to units from *in vivo* recordings and neurons from network models. To do so, we computed the lagged differences between spike times up to the 100^th^ order, i.e. the time differences between any spike and its successors (up to *n* = 100) at the unit level. The lagged differences were then sorted in 3.33ms bins from 0 to 1000ms. The resulting counts, once normalized, allowed to build the probability density function of the autocorrelogram, AC, which was smoothed by local non-linear regression (*loess* method, with span 0.1; to filter high frequency noise and correctly detect the peaks, see below) after removing its first 10ms, to eliminate source data contaminations, such as inter-spike intervals (ISIs) shorter than the absolute refractory period. We defined the peak of the autocorrelogram as its maximum, except when the maximum was the very first bin, in which case the peak was defined as the first local maximum after the first bin. The latency of the peak, LAT, was considered, for further analysis, as a structural parameter of the autocorrelogram characterizing the temporal signature of the neuron/unit spiking set. For each autocorrelogram, a global mono-exponential fit (GLOBAL fit) was then performed on the part of the autocorrelogram situated after the peak using the Levenberg-Marquardt algorithm (*nlsLM function in R*) for monkey data or von-Neumann–Karmarkar interior-point algorithm (*fmincon in Matlab*) for network models (we checked that either algorithm on the same spiking sets gave similar results), as following:

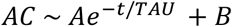

TAU, the time constant of the autocorrelogram fit characterized the temporal signature of the neuron., the amplitude of the exponential, and, the offset, are positive constants. Note that this mono-exponential fitting equation is strictly equivalent to that of Murray et al.(Murray et al., 2014), here corresponding to in the Murray method. Choosing one or the other did not affect the resulting fit and we kept the present form as it is easier to interpret. Fits on each autocorrelogram were performed 50 times, with random initial guesses in the range [0,2(*max(AC)* - *min(AC)*)] for *A*,[0,2*min(AC)*] for *B,* and [0,1000]ms for TAU, from which the best fit was kept.

In a minority of cases (less than 3% of neurons), the autocorrelogram following the peak (as defined above and denoted below the 1^st^ peak) could present a shape that diverged from a simple exponential decay, because of a fast and large dip, followed by a second local maximum, which preceded the slower, final exponential decay. In this case, we developed a pipeline aiming at consistently choosing the peak from which the fit started. To do so, we defined the autocorrelogram as having a dip if the first local minimum in the 100ms after the 1^st^ peak was below 75% of the global range of the autocorrelogram, *max(AC)* - *min(AC)*. In such cases, the second peak was defined as the maximum of the autocorrelogram after the dip and two additional mono-exponential fits of the autocorrelogram were performed, one from the first peak to the dip (FAST fit) and a second one from the second peak to the end of the autocorrelogram (SLOW fit). To be valid, any individual fit had to display positive *A*, *B* and TAU values. When neurons had a valid GLOBAL fit, two possibilities were considered. First, the valid GLOBAL fit was kept when at least one of the FAST and SLOW fits were not valid. Second, the valid GLOBAL fit was also kept when it was the best (i.e. its root-mean-square error was inferior to that of the sum of the valid FAST and SLOW fits) and excluded otherwise. Neurons that did not have a valid GLOBAL fit were also excluded from further analysis. Thus, while FAST and SLOW fits were *de facto* systematically excluded from further analysis, they were only used to ensure the quality of GLOBAL exponential fits. Note again that excluding less than 3% of neurons, this complex procedure was very conservative and designed for the sake of fitting performance.

### Hidden Markov Model (HMM) analysis

We used HMM to map the spiking set of neural network models and unit populations in monkeys onto discrete states of collective activity, based on previously established methods(Abeles et al., 1995; Seidemann et al., 1996). HMM methods allow to determine the probability *p*(*S*_*k*_(*t*)) of the network to be in state *S*_*k*_,*k* ∈{1…*n*_*s*_} at time *t* Typically, we found that, as previously shown in frontal areas, population activity organized into periods that lasted in the range ~10*ms* – 10*s*, i.e. transition probabilities were small and states were quasi-stationary. When all probabilities of being in a state *p*(*S*_*k*_) < 0.8, the network was considered to be in the null state *S*_0_, signifying that the network was not in any of the states. Periods in the *S*_0_ state were typically short (mean: LPFCm=16ms, MCCm=36ms, not shown). Thus, when immediately preceded and followed by two periods in the same state *S*_k_, periods in *S*_0_ were attributed the state *S*_k_. For each network spiking set assessed, we pooled the durations of all periods in all the states of the HMM model, to build the overall probability distribution of period durations *p(d)*. We then used this probability distribution to compute

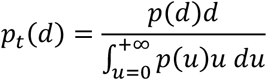

i.e. the proportion of time spent in state periods of duration *d*, that is, the probability, at any given instant in time, of being in state periods of duration *d*. We could not find any suitable method of stably determining the number of states *n*_*s*_. However, as a low number of states is more parsimonious in terms of data interpretation(Pohle et al., 2017) in general and because the task structure contains a low number of possible states in terms of actions (four), reward on the last trial (incorrect trial, first correct trial, correct trial after previous correct trials) and behavioral states (exploration, exploitation), we arbitrarily fixed *n*_*s*_ = 4. Each HMM model analysis was conducted on a spiking set lasting 600 s, both in neural network models and unit populations in monkeys. For each monkey area, the activity of all neurons was pooled, regardless of their recording session. This was mandatory because the number of neurons simultaneously recorded in each session was typically inferior to 5, so that HMM models were inefficient in detecting states. Pooling all neurons allowed the detection of global states that corresponded to the combination of collective dynamics recorded during distinct sessions, i.e. that were not time-locked together (phase information lost across sessions) and causally independent. Although chimeric, these HMM states were nevertheless able to indirectly capture the underlying temporal structure of collective spiking discharges in frontal areas in a similar way and thus allowed comparing LPFC and MCC collective temporal structure. In control HMM models, both the timing and neuron assignment of all spikes were randomly shuffled. The initial estimation of the average state duration across all periods in a given state was taken at a high value (300ms), which was suggested to give better log-likelihood scores and converge to similar states across repetitions of the HMM (Seidemann et al., 1996). The time bin was Δ*t* = 0.5*ms*

### Principal component analysis

The principal component analysis (PCA) of LPFC and MCC of monkeys’ units and neural network models’ neurons spiking activity was computed from firing frequencies, in order to better visualize and characterize collective dynamics. PCA was achieved on the set of the spiking frequency vectors of all units/neurons in each case. Spiking frequency was estimated through convolution of spiking activity with a normalized Gaussian kernel with standard deviation σ = 100*ms*, as average frequencies were typically < 10*Hz* in both areas. For each neuron, frequencies were then centered and standardized for optimal PCA. Cells with average frequencies less than 0.5 Hz were removed for the experimental data and for the model data, to avoid abnormal standardized frequencies when the neuron’s average frequency was too low (at most 6 cells per area).

### Perturbation protocol for state transitions

We assessed the contribution of excitatory and inhibitory neural populations to the stability of HMM states. To do so, we estimated the probability to stay in a given ongoing (or perturbed, see below) HMM state or to switch toward a distinct target (or perturbing) state in response to specified perturbations. The perturbation was achieved by substituting the value of neural variables (membrane potential, spiking state, calcium concentration, downstream channel opening probabilities) of a random subset of excitatory (respectively inhibitory) neurons of the ongoing state by those of the same neurons taken from the (distinct) target state. Specifically, starting from an initial (unperturbed) 600 s simulation, perturbations were achieved by substituting state variables 50ms after the onset of a randomly chosen period of a specified perturbed state by those taken 50ms after the onset of a randomly chosen period of a distinct perturbing state and the resulting network states used as initial conditions for further “perturbation simulations”. For each perturbation simulation, the network was simulated from the perturbation time to the end of the period when the network was not perturbed and the HMM state was determined as the posterior state probability based on HMM transition and emission matrices obtained from the entire initial unperturbed simulation. The probability to escape the ongoing state (Fig. 8.h, left) and to reach the target state (Fig. 8.h, right) were then computed as the proportion of time spent, during the ongoing period, in a HMM state different from the ongoing perturbed state (escape ongoing state probability), and in the target perturbing state (reach target state probability), respectively. The effects of perturbations were tested by replacing either excitatory or inhibitory populations, where proportions of replaced neurons systematically varied in the range 0-1. For each neuron type and proportion tested, the perturbation protocol was applied and results averaged for 50 random combinations of periods (with period durations > 100ms), for each of the 12 possible pairs of the 4 HMM states (excluding pairs of repeated states), over 20 different randomly initialized MCCs. Probabilities were offset and normalized to remove the basal probability of escaping the ongoing (0.09) and reaching the target (0.01) states when no perturbation was applied (such transitions were due to random selection of simultaneous spikes when initiating the HMM analysis).

### Behaviour and context-dependent modulations

#### Behavioural Task

Monkeys were trained to perform a dual task involving rule-based and internally driven decisions(Stoll et al., 2016). Monkeys performed the task using a touch screen. In each trial they could freely choose whether to perform a rewarded categorization task or to check their progress toward a large bonus juice reward (**Fig. 3a**). Upon checking (selection of a disk-shaped lever) progress was indicated by the onset of a visual ‘gauge’ (an evolving disk inside a fixed circle). Choosing the categorization task (selection of an inverted triangle lever) started a delayed response task in which an oriented white bar (cue) was briefly presented, followed by a delay at the end of which 2 bars oriented 45° leftward and rightward where presented. Selecting the bar matching the cue orientation led to a juice reward. An incorrect response led to no reward delivery. The gauge increased based on correct performance in the categorization task following 7 steps to reach the maximum size. If the animal checked while the gauge was full, the bonus reward was delivered, and the gauge reset to step 1. The full gauge was reached after either 14, 21, 28 or 35 correct trials (= number of trials to complete the 7 steps, pseudo-randomly chosen in each block). Thus, the gauge could increase at one of 4 different speeds.

#### Pause vs. engage periods

As each trial was self-initiated by the animal, monkeys could decide to take a break in their work. We defined pauses as periods of at least 60 seconds without trial initialization. Monkeys made on average 3.4±2.57 pauses per session (mean±sd, monkey A: 3.44±2.55, monkey H: 3.34±2.63; see **Fig. 3b**). We extracted spike times during the defined pause and engage time segments for each unit, and then extracted TAU using the method described above. We only kept units with successful TAU extraction for both periods (n_MCC-FS_=19, n_MCC-RS_=86, n_LPFC-FS_=29, n_LPFC-RS_ =95).

#### Fast vs. slow-paced blocks

We defined 14 and 21 correct trials blocks to be fast blocks and 28 and 35 correct trials blocks as slow blocks. We considered neuronal activity from the first-time monkeys checked in a block until the end of the block. We excluded pause periods from this analysis. We extracted spike timing from the segments and computed timescales as previously, keeping only units with successful timescale extraction for both periods (n_MCC-FS_=33, n_MCC-RS_=165, n_LPFC-FS_=46, n_LPFC-RS_ =165).

#### Emptier vs. fuller gauge size seen

In each block, monkeys used the gauge size observed upon checking to regulate their future decisions to check. The checking frequency increased with gauge size with a marked increase at steps > 4. We thus compared neuronal activity in periods in which monkeys saw gauges of size < 4, with periods in which they saw gauges > 4, excluding the very beginning of blocks when monkeys have not seen the gauge yet, and pauses periods. We perform this analysis on 430 units (n_MCC-FS_=30, n_MCC-RS_=178, n_LPFC-FS_=47, n_LPFC-RS_ =175).

To test whether current block speed had an influence on TAU at the unit level, we computed a modulation index for each unit: log(TAU_slow_)/log(TAU_fast_). Similarly, to test whether gauge filling state had an influence on TAU at the unit level, we computed a modulation index for each unit: log(TAU_empty_)/log(TAU_full_) where TAU_full_ corresponds to TAU calculated on the spike data recorded during the time in blocks where the gauge was superior of equal to the 4^th^ level.

### Statistical analyses

All analyses were performed using R (version 3.6.1) with the RStudio environment(R_core_team, 2014).

#### BLOM transformation

As some timescale measures are non-normally distributed, analyses required a robust non-parametric test. We opted for the BLOM transformation which is a subcase of Rank-Based Inverse Normal Transformations(Beasley et al., 2009). Basically, the data is ranked and then back transformed to approximate the expected normal scores of the normal distribution according to the formula:

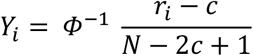

where r_*i*_ is the ordinary rank and Y_*i*_ the BLOM transformed value of the *i*th case among the N observations. Φ-1 is the standard normal quantile (or probit) function and c a constant set to 3/8 according to Blom(Blom, 1958). Regular parametric analyses can then be performed on the transformed data. Since z-scores of the transformed data are normally distributed and differences are expressed in standard errors, main effects and interactions can easily and robustly be interpreted. As sanity checks we also ran more classical non-parametric tests (Wilcoxon test) on non-normally distributed data leading to the same conclusions.

### Task-related analyses

#### Single unit activity

Each unit’s spikes were counted in sliding bins of 200ms overlapping by 50ms from feedback onset to 800ms post-feedback and during the intertrial interval from 400ms before the end of trial signal onset to 2000ms after its onset.

#### Group analyses using a glmm

We used a *glmm* using a Poisson family. p-values were corrected for multi-comparison with the false discovery rate algorithm with the number of comparisons being the number of timebins (p.adjust function in R).

The mixed models used were of the form:

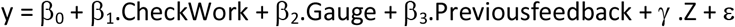

where γ.Z is the random term, and CheckWork, Gauge and PreviousFeedback are the fixed effects describing the Check versus Work decision (0/1), the gauge size (1–7) and the feedback in the previous trial (0/1) with their respective parameters (β). In the *glmm*, the Single unit identity was used as a random factor.

A persistent problem with Poisson models in biology is that they often exhibit overdispersion. Not accounting for overdispersion can lead to biased parameter estimates. To deal with overdispersion we used observation-level random effects (OLRE), which model the extra variation in the response variable using a random effect with a unique level for every data point.

#### Median splits

To test the hypothesis that units with different timescales may encode feedback differently we divided the units into two groups based on the median of the timescale metric. We computed the median of the metric (e.g. peak latency or TAU) in all the units of a given cell type. Then we put units with a metric value below the median into the ‘short’ group and units with a metric value above the median into the ‘long’ group.

### Timescale and coding variations along the antero-posterior axis

We considered the genu of the arcuate sulcus as an anatomical landmark from which we computed distances of recording location along the anterior-posterior axis from MRI reconstructions. We questioned TAU antero-posterior variability keeping recording locations covering the same range in both monkeys. We ordered locations from the most posterior site for each area. We excluded FS units from statistical analysis due to their disparateness (RS units, monkey A: n_MCC_=112, n_LPFC_=110; monkey H: n_MCC_=54, n_LPFC_=64). This analysis was conducted separately between monkeys to account for inter-subject anatomical variability.

To test variation in population coding along the antero-posterior axis we divided single-units into a posterior and anterior group based on the range of locations of each area (MCCpost from 4.5 to 7mm, n_MCCRSpost_=84, n_MCCFSpost_=14; MCCant from 7 to 9.5mm, n_MCCRSant_=82, n_MCCFSpost_=16; LPFCpost from 2.5 to 6mm, n_LPFCRSpost_=77, n_LPFCFSpost_=19; LPFCant from 6 to 8.5mm, n_LPFCRSant_=97, n_LPFCFSant_=19). Population coding analysis is described in **Task-related analyses**.

### Cellular model of pyramidal neurons in frontal areas

We built a generic biophysical Hodgkin-Huxley model of the detailed dynamics of membrane potential and of ionic and synaptic currents of individual pyramidal neurons in frontal areas. The model was generic, being endowed with a large set of ionic voltage- and calcium-dependent conductances, to encompass the wide possible repertoire of spiking discharge patterning encountered *in vivo*. In the model, the membrane potential followed

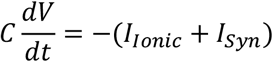

where C is the specific membrane capacity and the membrane ionic current writes

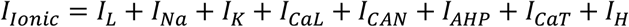

in which the leak current is

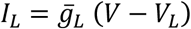

and action potential (AP) currents (*I*_*Na*_, *I*_*K*_) are taken from a previous model we devised to reproduce spike currents of frontal pyramidal regular-spiking neurons(Naudé et al., 2012). The high-threshold calcium current was

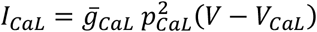

where the activation followed first-order kinetics

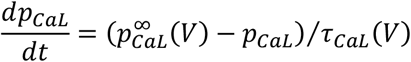

with a voltage-dependent time constant

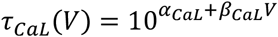

where α_CaL_ and β_CaL_ were fitted from *in vitro* data(Helton et al., 2005). The infinite activation followed

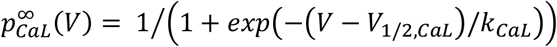

where V_1/2,CaL_ and *k*_*CaL*_ respectively denote the half-activation potential and e-fold slope of the Boltzmann activation voltage-dependence, estimated from *in vitro* data(Helton et al., 2005).The cationic non-selective (*I*_*CAN*_) current and the medium after-hyperpolarization (*I*_*AHP*_) current, responsible for frequency adaptation in pyramidal neurons were taken as in Rodriguez et al., 2018, with

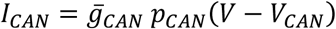

and

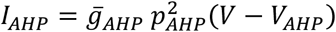

The activation of both currents, *p*_*x*_(x ∈ {*CAN, AHP*}) followed

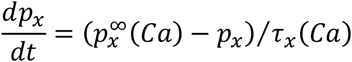

with

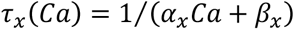

and

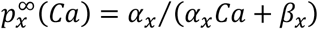

where α_*x*_ and β_*x*_ respectively denote activation and deactivation kinetic constants consistent with experimental data in layer 5 PFC pyramidal neurons(Faber and Sah, 2007; Haj-Dahmane and Andrade, 1997; Villalobos et al., 2004). The low-threshold calcium (*I*_*CaT*_) and hyperpolarization-activated (*I*_*H*_) currents were from reference(Ritter-Makinson et al., 2019). To account for autocorrelogram parameters, we employed different versions of the model that contained distinct subsets of ionic currents, which have been implicated in adaptation and bursting (*I*_*CaL*_, *I*_*AHP*_), rebound (*I*_*CaT*_, *I*_*H*_), and regenerative and bistable discharge (*I*_*CaL*_,*I*_*CAN*_,*I*_*AHP*_) in cortical pyramidal neurons (see *Results* and *Supplementary Material*). Calcium concentration dynamics resulted from the inward influx due to and and first-order buffering or extrusion(Rodriguez et al., 2018) through:

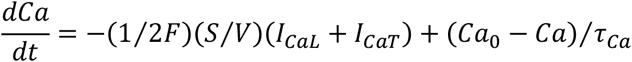

where is the Faraday constant, Ca_0_ is the basal intracellular calcium concentration, τ_Ca_ is the buffering time constant, and

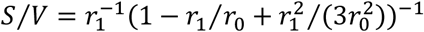

is the surface area to volume ratio of an idealized intracellular shell compartment of thickness situated beneath the surface of a spherical neuron soma of radius r_0_.

The synaptic current (*I*_*Syn*_) mimicked *in vivo* conditions encountered by neurons in the asynchronous irregular regime, summing random synaptic excitatory inputs, through AMPA and NMDA receptors, and inhibitory inputs, through GABA_A_ and GABA_B_ receptors. Thus,

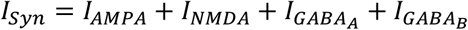

For AMPA, GABA_A_ and GABA_B_,

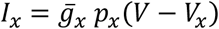

where *p*_*x*_ is the opening probability of channel-receptors and *V*_*x*_ the reversal potential of the current. The NMDA current followed

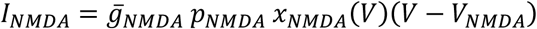

incorporating the magnesium block voltage-dependence modeled(Jahr and Stevens, 1990) as

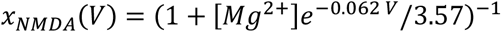

To simulate fluctuations encountered *in vivo*, all opening probabilities followed Ornstein-Uhlenbeck processes(Destexhe and Paré, 1999)

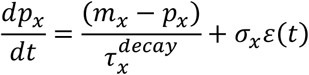

where *ε*(*t*) is a Gaussian stochastic process with zero mean and unit standard deviation and *m*_*x*_ and σ_*x*_ are the mean and standard deviation of the opening probabilities. For AMPA and GABA_A_, the mean was taken as the steady-state value of first-order synaptic dynamics described in the network model (see below):

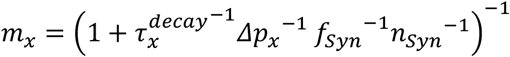

with *n*_*syn*_ pre-synaptic neurons firing at a frequency *f*_*syn*_ (with *Syn* ∈ {*Exc*, *Inh*} depending on the type of current considered), an instantaneous increase Δ*p*_*x*_ of opening probability upon each pre-synaptic spike and first-order decay dynamics with time constant 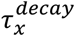 between spikes. For NMDA and GABA_B_, the mean was taken as the steady-state value of second-order synaptic dynamics described in the network model (see below):

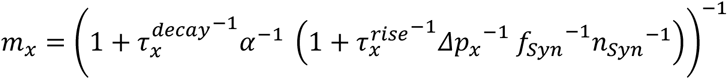

For all currents, standard deviations were taken as σ_*x*_ = 0.5*m*_*x*_. Feed-forward excitatory and inhibitory currents were balanced (Xue et al., 2014), according to the driving forces and the excitation/inhibition ratio, through

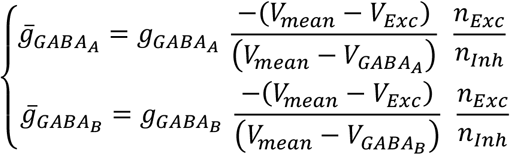

### Model of local recurrent neural networks in frontal areas

We built a biophysical model of a generic local frontal recurrent neural network, endowed with detailed biological properties of its neurons and connections. The network model contained neurons that were either excitatory (E) or inhibitory (I) (neurons projecting only glutamate or GABA, respectively(Dale, 1935)), with probabilities *p*_*E*_ and *p*_*I*_ = 1 − *p*_*E*_ respectively, and *p*_*E*_/*p*_*I*_ = 4(Beaulieu et al., 1992). Connectivity was sparse (i.e. only a fraction of all possible connections existstho(Thomson, 2002)) with no autapses (self-connections) and EE connections (from E to E neurons) drawn to insure the over-representation of bidirectional connections in cortical networks (four times more than randomly drawn according to a Bernoulli scheme(Song et al., 2005)). The synaptic weights *w*_*(i,j)*_ of existent connections were drawn identically and independently from a log-normal distribution of parameters μ_*w*_ and σ_*w*_ (Song et al., 2005). To cope with simulation times required for the massive explorations ran in the parameter space, neurons were modeled as leaky integrate-and-fire (LIF) neurons, i.e. the AP mechanism was simplified, compared to the cellular model (see above). Moreover, leveraging simulations at the cellular level, we only considered the *I*_*CAN*_ and *I*_*AHP*_ amongst the ionic currents of the cellular model (see above). Thus, the membrane potential followed

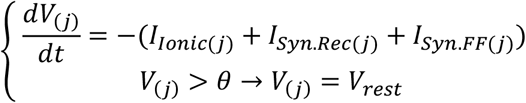

where repolarization occurred after a refractory period Δ*t*_*AP*_. The ionic current followed

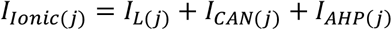

with parameters and gating dynamics of ionic currents identical to the cellular model. The intra-somatic calcium concentration *Ca* evolved according to discrete spike-induced increments and first-order exponential decay:

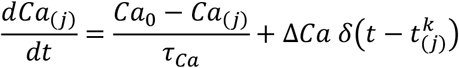

where 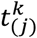 is the time of the *k*_th_ spike in the spike train of neuron *j*, *δ* the Dirac delta function, τ_*ca*_ the time constant of calcium extrusion, *Ca*_0_ the basal calcium and Δ*Ca* a spike-induced increment of calcium concentration. The recurrent synaptic current on post-synaptic neuron *j*, from – either excitatory or inhibitory – pre-synaptic neurons (indexed by *i*), was

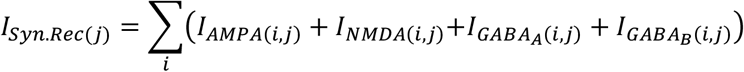

The delay for synaptic conduction and transmission, Δ*t*_syn_, was considered uniform across the network(Brunel and Wang, 2001). Synaptic recurrent currents followed

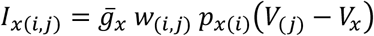

where *w*_*(i,j)*_ is the synaptic weight, *p*_*x(i)*_ the opening probability of channel-receptors and V_X_ the reversal potential of the current. The NMDA current followed

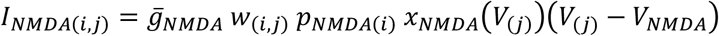

with *x*_*NMDA*_(*V*) the magnesium block voltage-dependence (see cellular model). AMPA and GABA_A_ rise times were approximated as instantaneous (Brunel and Wang, 2001) and bounded, with first-order decay

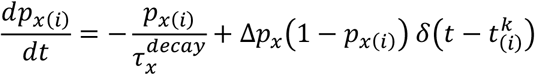

To take into account the longer NMDA (Wang et al., 2008) and GABA-B (Destexhe et al., 1998) rise times, opening probabilities followed second-order dynamics (Brunel and Wang, 2001)

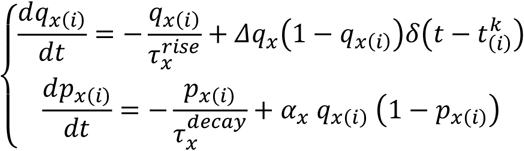

Recurrent excitatory and inhibitory currents were balanced in each post-synaptic neuron (Xue et al., 2014), according to driving forces and excitation/inhibition weight ratio, through

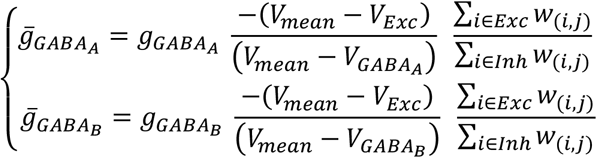

with *V*_*mean*_ = (θ + *V*_*rest*_/2 approximating the average membrane potential.

The feed-forward synaptic current *I*_*Syn.FF(j)*_ (putatively arising from cortical and sub-cortical inputs) consisted of an AMPA component

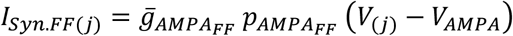

with a constant opening probability 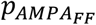.

### Numerical integration and parameters of the models

Models were simulated and explored using custom developed code under MATLAB and were numerically integrated using the forward Euler method with time-steps Δ*t* = 0.1*ms* in cellular models and Δ*t* = 0.5*ms* in network models.

Unless indicated in figure legends, standard cellular parameter values were as following. Concerning ionic currents
*C* = 1*μF.cm*^−2^, 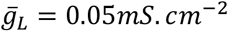, *V*_*L*_ = −70*mV*, 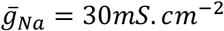, *V*_*Na*_ = 50*mV*, 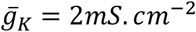, *V*_*K*_ = −90*mV*, 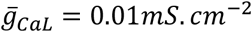, *V*_*CaL*_ = 150*mV*, 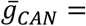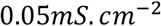, *V*_*CAN*_ = 30*mV*, α_*CAN*_ = 0.0015μM^−1^.ms^−1^, β_*CAN*_ = 0.005*ms*^−1^, 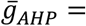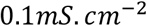, *V*_*AHP*_ = −90*mV*, α_*AHP*_ = 0.025*μM*^−1^, β_*AHP*_ = 0.025*ms*^−1^, 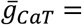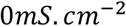, *V*_*CaT*_ = 120*mV*, 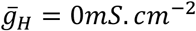, *V*_*H*_ = −40*mV*, V_*τH*1/2_ = −105*mV*, *k*_*τH*_ = 10*mV*, τ_*H,min*_ = 1000*ms*, τ_*H,max*_ = 6000*ms*, *Ca*_0_ = 0.1*μM*, τ_*ca*_ = 0.1*μM*, τ_*ca*_ = 25*ms*, *F* = 96500 *mol.s*^−1^.A^−1^, r_0_ = 4 ·10^−4^*cm*, r_1_ = 0.25 ·10^−4^*cm*. Concerning synaptic currents, 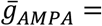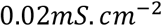, 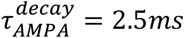, 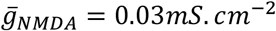, α_*NMDA*_ = 0.275*ms*^−1^, 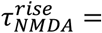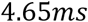, 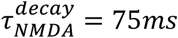, *V*_*AMPA*_ = *V*_*NMDA*_ = 0*mV*, 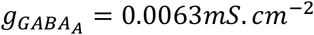, 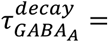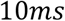, 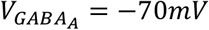, 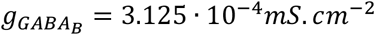, 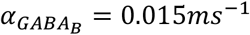, 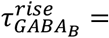, 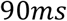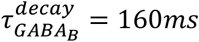, 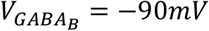, 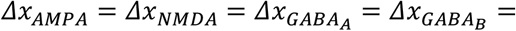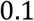, *V*_*mean*_ = −57.5*mV*, *n*_*Exc*_ = 484, n_*Inh*_ = *n*_*Exc*_/4 = 121, *f*_*Exc*_ = *7Hz*, *f*_*Inh*_ = *7Hz*, [*Mg*^2+^] = 1.5*mM*.

Unless indicated in figure legends, standard parameter values in network models were identical to cellular model parameters, except for the following. Concerning the network, *N* = *n*_*Exc*_ + *n*_*Inh*_ = 605 neurons, *p*_*Exc*_ = 0.8, so that *n*_*Exc*_ = *Np*_*Exc*_ = 484 and *n*_*Inh*_ = *Np*_*Inh*_ = 121. Concerning the weight matrix, *μ*_*w*_ = 0.03, *p*_*EE*_ = *p*_*EI*_ = *p*_*II*_ = 0.3, *p*_*IE*_ = 0.55. Concerning Integrate-and-Fire neuron properties and intrinsic currents, *V*_*rest*_ = −65*mV*, *θ* = −50*mV*, *V*_*mean*_ = (*V*_*rest*_ + θ)/2 = −57.5*mV*, *Δt*_*AP*_ = 3*ms*, *ΔCa* = 0.2*μM*, 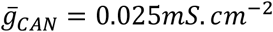.Concerning synaptic currents, *Δt*_*syn*_ = 0.5*ms*, 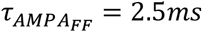, 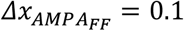, 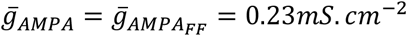, 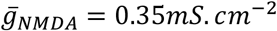, 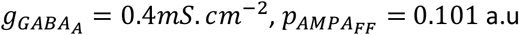.

### Model similarity tomonkey data

The bivariate probability density distribution of neuronal TAU and LAT autocorrelogram parameters was estimated in RS and FS units in monkey in both the LPFC and MCC, using bivariate normal kernel density functions. For cellular models, similarity maps to monkey data was determined as following: for each model parameter couple of the map, the similarity to the considered cortex (PFC or MCC) was defined as the probability density of that cortex to display the TAU and LAT parameters produced by the model. Cellular models with mean firing frequency superior to 20 Hz were considered to discharge in an unrealistic fashion, compared to data, and were discarded. In network models, for each parameter value (one-dimensional explorations) or model parameter couples of the map (two-dimensional explorations), the similarity (S) was defined as the normalized Frobenius inner product between the bivariate probability density distributions of units in monkeys (U) and that of neurons in the network model (N), following

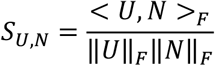

In order to account for the TAU and LAT autocorrelogram parameters for both RS and FS populations, the similarity was calculated separately as RS with Exc and FS with Inh. Seeing as excitatory neurons represent *p*_*Exc*_ = 0.8 of the neurons in cortex (Beaulieu et al., 1992), the overall similarity was then calculated as

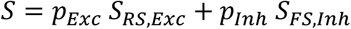

## Additional information

## Acknowledgement

This work was supported by the Medical Research Foundation (FRM) (*Equipe* DEQ20160334905, and VF: FDT201904008187), the French National Research Agency (DECCA ANR-10-SVSE4-1441 and PARABIST ANR-16-NEUC-0006-01), and was performed within the framework of the labex CORTEX ANR-11-LABX-0042 of Université de Lyon. EP is employed by the Centre National de la Recherche Scientifique. We thank C. Nay for administrative support and C. Wilson and J. Naudé for helpful discussions and proofreading the manuscript.

## Author contributions

VF, FS and EP acquired and analyzed experimental data; VF, MS, BD and EP designed the analysis procedures; MS and BD designed, constrained, simulated models and analyzed modelling results; VF, FS, MS, BD and EP wrote the article.

## Data availability

All spike time series from monkey recordings and scripts for temporal signatures extraction are freely accessible on the zenodo repository (https://doi.org/10.5281/zenodo.3947745). Scripts of computational models are available upon request.

