## Supplementary materials for "Inhibitory control of frontal metastability sets the temporal signature of cognition"

### Supplementary material

**Timescales distributions considering 3 populations of cell types based on spike widths.**
We clustered units in cell populations based on spike waveform features, notably spike width (**Fig. 1c** in main text). In the main analysis we pooled together units with large and very large spike widths into a Regular Spiking population (RS, long spikes; nMCC=257, nLPFC=215 units) that we contrasted with Fast Spiking cells (FS, short spikes; nMCC=37, nLPFC=61 units) for timescale descriptions. We verify here that considering 3 cell populations (FS, short spikes: nMCC=37, nLPFC=61 units; RS1, long spikes: nMCC=183, nLPFC=162 units and RS2 longer spikes: nMCC=36, nLPFC=28 units) lead to the same conclusions than when pooling the two RS populations. As in Fig. 2c, TAU are higher in the MCC than in the LPFC (**Fig. S1a**, linear model fit on BLOM transformed TAU for normality, TAU = Area * Unit type, $F(5,487)=28.4$, Area : $p<10^{-4}$, Unit type: $p=0.05$, interaction: $p=0.91$). Autocorrelogram peak latencies were longer for both MCC RS1 and RS2 populations (**Fig. S1b**, linear model fit on BLOM transformed Latency for normality, Latency = Area * Unit type, $F(5,487)=23.63$, interaction: $p<0.005$).

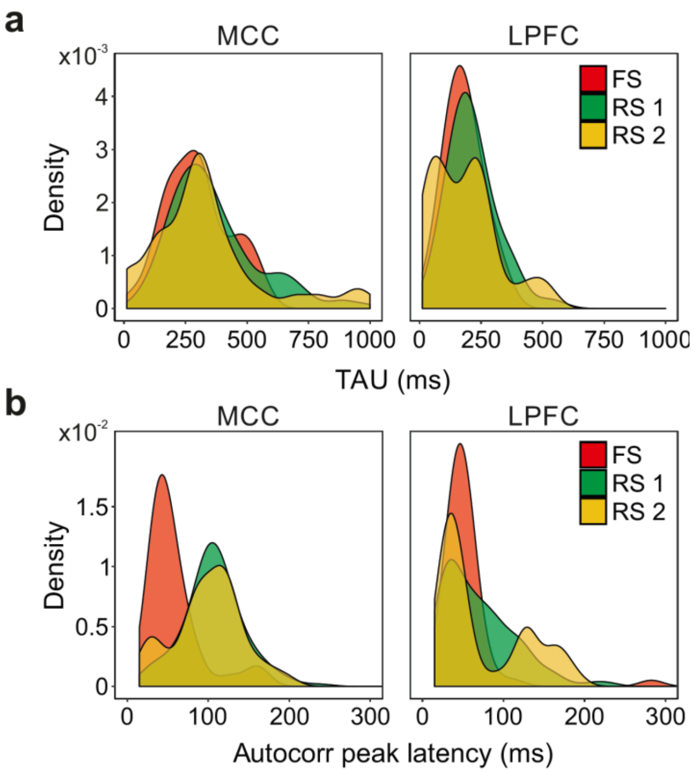


**Figure S1.** Spike autocorrelogram features considering 3 populations in the MCC and LPFC. (**a**) TAUs and (**b**) LAT distribution in the 3 cell populations in the MCC and the LPFC. Densities were computed using a gaussian kernel with standard deviation of 50ms and 15ms respectively for TAU and peak latencies.

**Relationship between firing rate and temporal signatures**

As both the firing rate and TAU are features extracted from the distribution in time of action potentials, the relationship between the two remained a persistent question throughout our analyses. Firing rates are classically computed in bins of hundreds of milliseconds around events of interest to capture the activity dynamics. In the context of our study which describes intrinsic properties of units, we think that the maximum firing rate of a given units is informative of its physiological range of activity. We computed the firing rate of units in 2 second bins and then considered the maximum firing rate as the average of the bins with firing rate above the third quartile. Varying the bin width (from 200ms to 2sec by step of 200ms) did not change the firing rate measure extracted. The population of FS has higher firing frequency (Supplementary Fig. 2a; linear model fit on BLOM transformed firing rate for normality, FiringRate = Area * Unit type, $F(3,566)=12.61$, $p<10^{-7}$, interaction: ns, Area: ns, Unit type: $p<10^{-4}$). There is no relationship between the firing rate and TAU (Supplementary Fig. 2b; Pearson's correlation: $r(491)=0.060$, $p=0.19$).

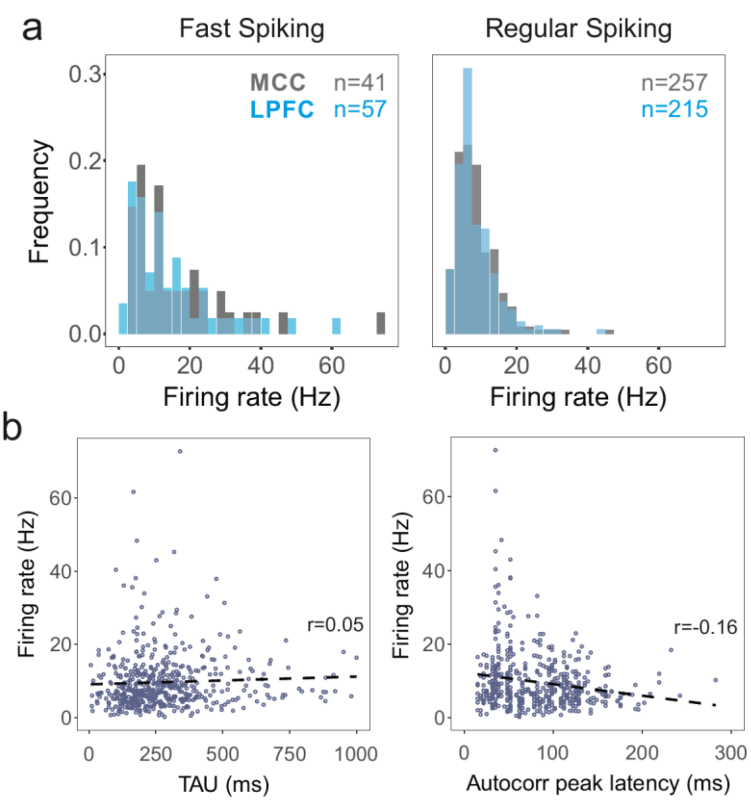


**Figure S2a.** Relationship between firing rate and timescale (**a**) Distributions of firing rates for fast spiking and regular spiking units in MCC and LPFC. (**b**) Scatter plot of firing rate as a function of TAU. There is no significant correlation between the two.

**Anatomical organization of temporal signatures**

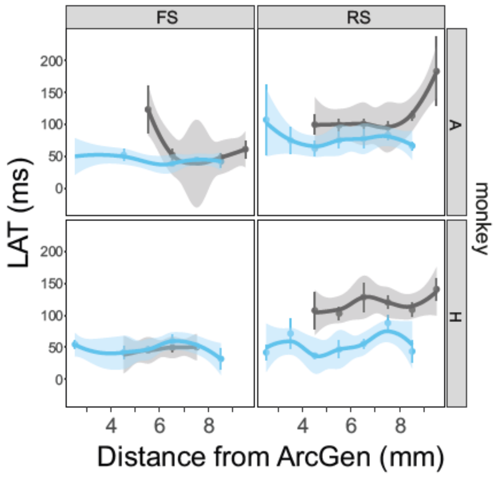


**Figure S2b. Anatomical distribution of LAT in MCC and LPFC for the 2 monkeys.**

We tested whether units with different temporal signatures were topographically organized across the recording sites. Because of the limited extent of positions on the mediolateral axis, and because of previous suggestions of organization along the anteroposterior axis[1,2], we focused on the latter. TAU and LAT averaged over neurons were computed for each level. For the two animals, coordinates were referenced to the anterior level of the genu of the arcuate sulcus (ArcGen)[3]. We tested whether TAU and LAT differed depending on the position of units in the rostro-caudal axis. TAU and LAT were not homogeneous along the anteroposterior axis. Specific monkey per monkey analyses specifically for TAU of RS units are presented in the main text. Data for LAT are presented in **Fig. S2b**. Statistics showed no significant variation of LAT along the MCC rostro-caudal axis, and a significant variation for monkey H in LPFC (ANOVA on Blom transformed LAT: MCC, monkey A: $F(5,112)=1.4$, $p=0.47$, monkey H: $F(5,54)=0.6$, $p=1$, LPFC, monkey A: $F(6,110)=1.0$, $p=0.86$, monkey H: $F(6,64)=2.7$, $p=0.046$). Linear regression analyses revealed only a positive trend in the MCC for monkey A (linear regression on Blom transformed LAT: MCC, monkey A: $t(1,112)=5.1$, $p=0.05$, monkey H: $t(1,54)=1.2$, $p=0.54$, LPFC, monkey A: $t(1,110)=1.4$, $p=0.49$, monkey H: $t(1,64)=4.0$, $p=0.10$ ; all p-values are FDR corrected for n=2 comparison per monkey).

**Parametric explorations in the pyramidal neuron model**

Biophysical properties of neurons can affect autocorrelation parameters in several ways. In principle, increasing the refractory period (through increased hyperpolarizing ionic conductance) shifts the distribution of 1st order lags (ISIs), thus increasing LAT (**Fig. S3**a i)[4]. Increasing burstiness of the spike discharge (through increased depolarizing conductance-mediated positive feedback) also increases the latency, because higher-order lag distributions are more peaked. Moreover, conductances with slow time constants (including many bursting-mediating conductances) increase that of the autocorrelogram itself. Finally, all these factors may interact in complex ways *in vivo* to set the spiking pattern that shapes autocorrelations.

We first explored these alternatives with a detailed biophysical Hodgkin-Huxley model of a generic frontal pyramidal cortical neuron, simulated in *in vivo* conditions. Pyramidal neurons display a huge electrophysiological diversity set by ionic channels, which, together with synaptic inputs, influences spiking patterns. Two conductances, i.e. cationic non-specific (CAN) and potassium after-hyperpolarization (AHP), were the sole couple able to affect both the LAT and TAU of the autocorrelation (compare **Fig. S3a** ii & iii). Interestingly, these conductances are prominent in monkey LPFC and MCC, as well as rodent prefrontal pyramidal neurons where they control regenerative discharge, bistability and burstiness[5–9]. Within physiological ranges, 1) the autocorrelogram LAT essentially increased with the maximal $g_{AHP}$ conductance, while 2) TAU increased in an intermediate range of $g_{AHP}$ and increased with $g_{CAN}$ (**Fig. S3b**), possibly accounting for differences between LPFC and MCC in monkeys. Remarkably, the low-threshold calcium (CaT), high-threshold calcium (CaL) and hyperpolarization-activated H conductances, which are ubiquitous and govern spiking patterns through spiking adaptation and rebound, as well as NMDA and GABA-B synaptic input conductances, which display long time constants, were all ineffective in adequately modulating autocorrelation parameters (**Fig. S4**).

Computing an estimation of the bivariate probability density distribution of neuronal autocorrelogram parameters for LPFC and MCC RS units (**Fig. S3c**) allowed to build a map of the similarity of the cellular model to RS units temporal signatures in monkey LPFC and MCC, defined as the bivariate probability density observed for the LAT and TAU yielded by the cellular model, given a ($g_{CAN}$, $g_{AHP}$) couple of parameters (see *Online Methods*). We found that the model displayed large (i.e. sub-maximal) similarity to the LPFC in a substantial region of ($g_{CAN}$, $g_{AHP}$) parameters (**Fig. S3d**). By contrast, this was not true for the MCC (**Fig. S3d**), because the model was unable to generate LAT in the 100-150 ms range that characterizes the MCC (**Fig. S3b**), even when exploring large ranges of CAN and AHP conductance kinetic parameters.

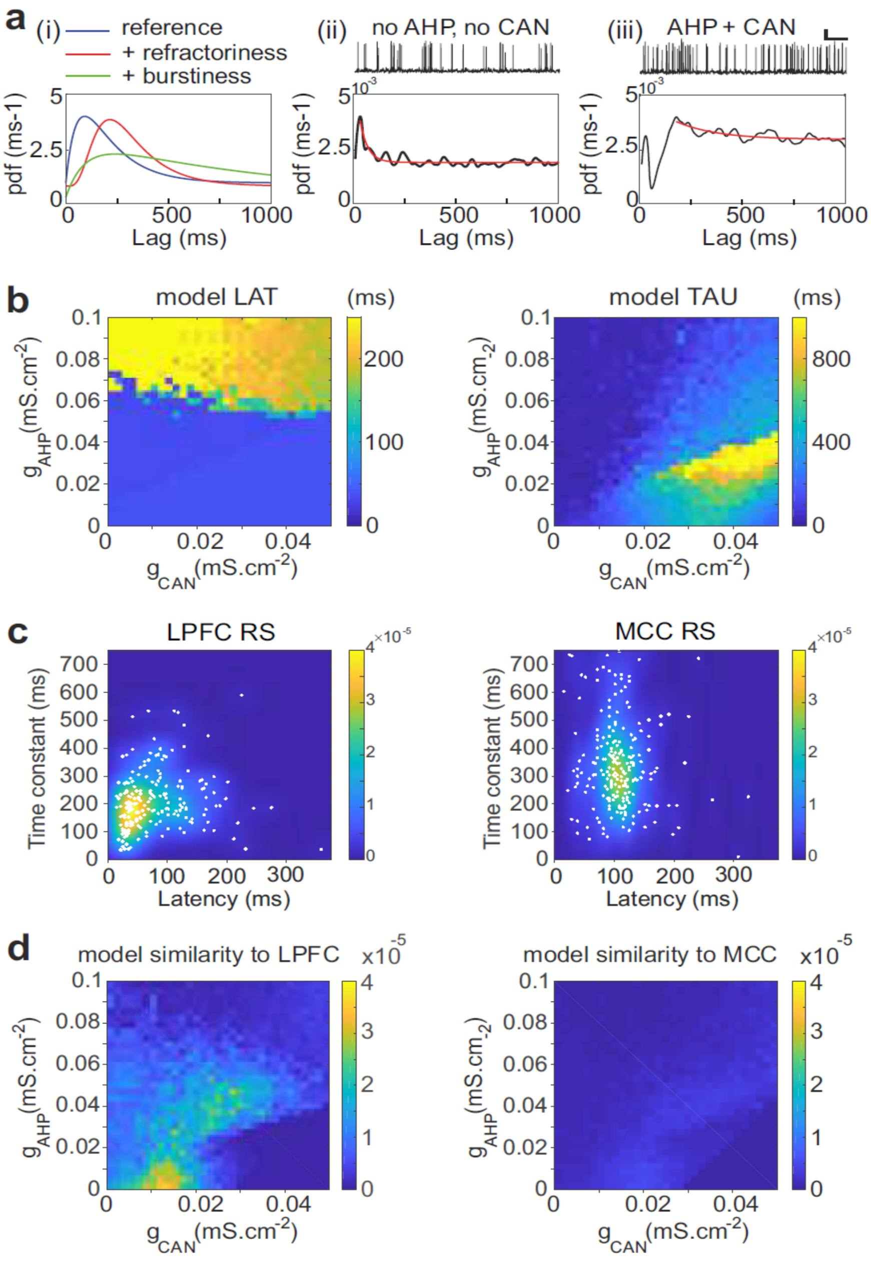


**Figure S3. Temporal signature in the pyramidal biophysical neuron model.** (**a**) In (i) schematic shapes of the autocorrelogram of a neuron (e.g. regular spiking, blue), of the same neuron with a larger refractory period (e.g. due to increased hyperpolarizing ionic conductance, red) or with higher burstiness of the spike discharge (e.g. due to increased depolarizing conductance-mediated positive feedbacks, green). (ii-iii) Autocorrelogram of a model pyramidal neuron (ii) in the absence ($g_{CAN}$=0mScm$^{-2}$, $g_{AHP}$=0mScm$^{-2}$) of CAN or AHP, or (iii) with $g_{CAN}$=0.05mScm$^{-2}$ and $g_{AHP}$=0.1mScm$^{-2}$. Scaling bars 1s and 25 mV. (**b**) Maps of the autocorrelogram latency (left) and time constant (right), as a function of $g_{CAN}$ and $g_{AHP}$ maximal conductances. (**c**) Bivariate probability density distribution of neuronal autocorrelogram LAT and TAU in RS units in both the LPFC (left) and MCC (right) in monkeys. (**d**) Similarity of the temporal signature between the frontal pyramidal neuron model and the population of RS units in the LPFC (left) and MCC (right), as a function of $g_{CAN}$ and $g_{AHP}$ maximal conductances.

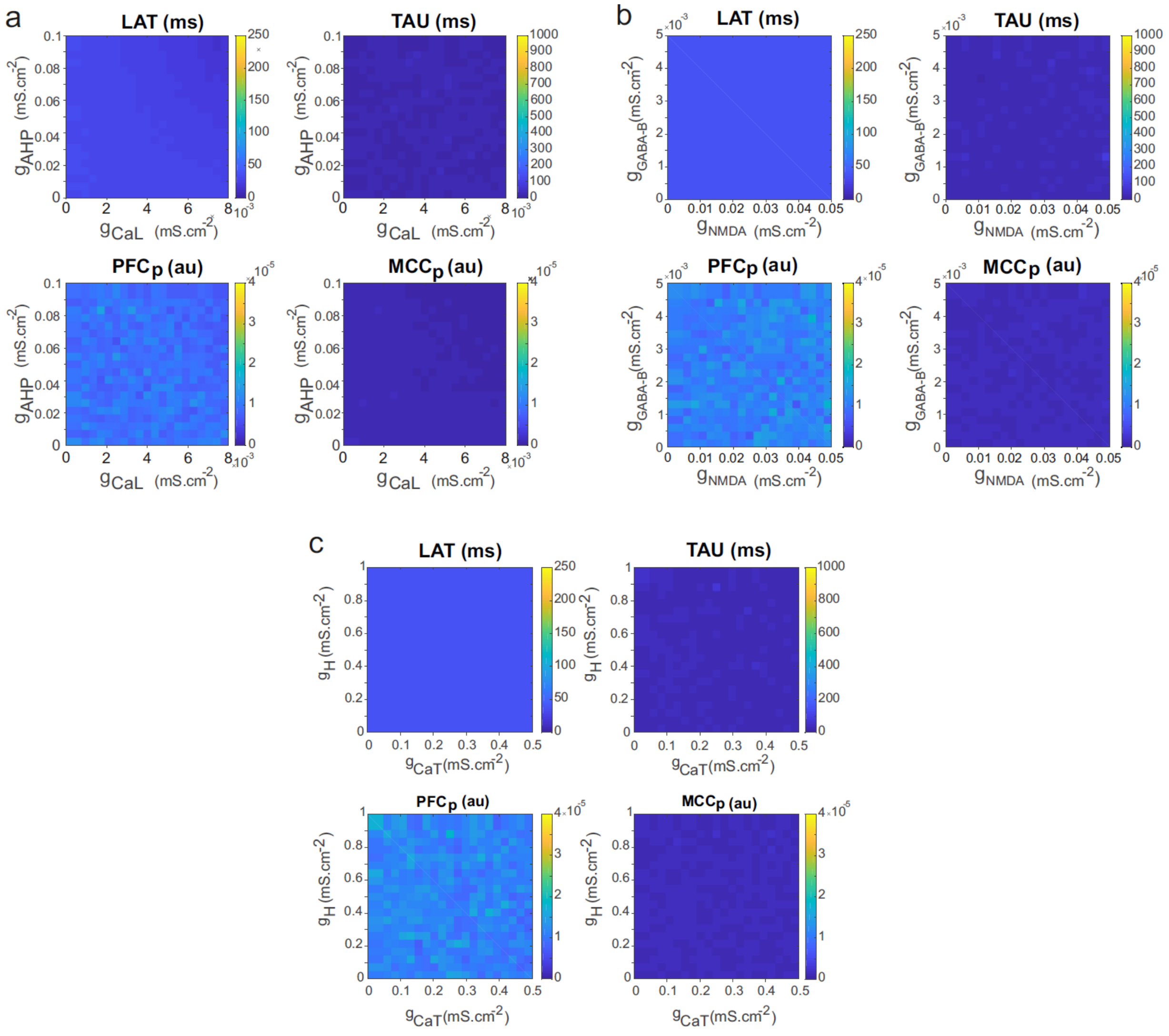


**Figure S4. Temporal signature in the pyramidal neuron model as a function of adaptation and rebound intrinsic and slow synaptic conductances.** Maps of LAT (left) and TAU (right), as a function of (**a**) the high-threshold calcium and after-hyperpolarization maximal conductances ($g_{CaL}$ and $g_{AHP}$) that set adaptation properties, (**b**) the low-threshold calcium (CaT) and hyperpolarization-activated H maximal conductances ($g_{CaT}$ and $g_{H}$) that set post-inhibitory rebound properties, and (**c**) the synaptic NMDA and GABA-B maximal conductances ($g_{NMDA}$, $g_{GABA\text{-}B}$) that set slow synaptic transmission.

**Parametric explorations in the network model**

We first assessed whether variations of a single biophysical parameter could explain the differences in the temporal signature of the MCC, compared to the LPFC: an increased TAU for RS and FS units and an increased LAT for RS (but not for FS) units. To do so, we tested many biophysical parameters determining the architectural, synaptic and intrinsic properties of the network, but none were able to account for these differences between frontal areas (not shown).

However, these explorations unraveled four model parameters of interest that were able, when varied within their physiological range (i.e. realistic regimes of network activity), to 1) affect either LAT in Exc (but not Inh) neurons, TAU in Exc neurons or TAU in Inh neurons, and 2) do so in a gradual fashion, i.e. allowing some possible form of (developmental, homeostatic or plastic) regulatory control. Indeed, several other parameters could vary TAU or LAT, but they did so abruptly, because their effects occurred at the vicinity of network bifurcations, where network activity dramatically saturated or was silenced, i.e. in non-physiological regimes.

The parameters of interest were the maximal conductances, on the one-hand, of two membrane ionic currents setting the intrinsic excitability and spiking pattern of cortical pyramidal neurons ($g_{CAN}$ and $g_{AHP}$) and, on the other hand, of two neurotransmitter-gated channels that set slow synaptic neurotransmission in cortical networks ($g_{NMDA}$ and $g_{GABA\text{-}B}$).

Firstly, decreasing $g_{CAN}$, the maximal conductance of the spike-triggered calcium-dependent cationic current, which controls regenerative discharge and spiking bistability in pyramidal neurons[6], gradually increased LAT in Exc neurons, although this occurred at low values where the network was nearby silence and displayed very low firing frequency (**Fig. S5a, upper left).** The CAN current is absent in Inh neurons[10], so that changing $g_{CAN}$ (i.e. only in Exc neurons) left LAT constant in Inh neurons (**Fig. S5a, lower left**). Besides, increasing the CAN maximal conductance gradually increased TAU in Exc neurons (**Fig. S5a, upper right**). This arose because $g_{CAN}$ increases burstiness, a factor that can increase the autocorrelogram time-constant[4]. However, it had no effect on TAU in Inh neurons, where it is absent (**Fig. S5a, lower right**). Thus, while $g_{CAN}$ possibly accounted for LAT in frontal areas (increased LAT in MCC Exc neurons, no change in Inh neurons), as well as for the increased TAU in MCC Exc neurons, it could not explain the TAU difference in Inh neurons (**Table S1**). Moreover, accounting for LAT and TAU in the MCC required incompatible $g_{CAN}$ ranges of values and this was the same for LPFC (**Table S1**).

Secondly, the maximal conductance of the medium AHP current ($g_{AHP}$), a spike-triggered calcium-dependent potassium current, which balances the CAN current in the patterning of spiking in pyramidal neurons, increased LAT in Exc neurons (**Fig. S5b, upper left**); this arose because AHP increases the refractory period, which can increase LAT[4]. Similarly to CAN, the AHP current is absent in Inh neurons[10], so changing $g_{AHP}$ (i.e. in Exc neurons) left LAT constant in Inh neurons (**Fig. S5b, lower left**). Besides, although $g_{AHP}$ is largely known for its effect on firing frequency adaptation, an important determinant of discharge temporal patterning, it displayed an extremely weak effect on TAU (**Fig. S5b, right**). Thus, while $g_{AHP}$ possibly accounted for LAT in frontal areas (increased LAT in MCC Exc neurons, no difference in Inh neurons), it could not explain differences in TAU (**Table S1**).

Together, these effects of intrinsic conductances at the network scale shared important trends with those in the cellular model, inasmuch as $g_{AHP}$ increased LAT and $g_{CAN}$ increased TAU in Exc neurons.

Thirdly, the NMDA receptor maximal conductance, $g_{NMDA}$, displayed no effect on LAT (**Fig. S5c, left**), but it increased TAU (**Fig. S5c, right**) in Exc and, to a lesser extent, in Inh neurons, because of its slow synaptic action (decay time constant, 75ms) on both neuronal types. However, these effects on TAU occurred at high $g_{NMDA}$ values where the network was nearby saturation and displayed unrealistic high frequency activity. Thus, while $g_{NMDA}$ possibly accounted for TAU in frontal areas and for the absence of change in LAT in Inh neurons, it could not explain the difference in LAT in Exc neurons between LPFC and MCC (**Table S1**).

Fourthly, the $GABA_B$ receptor maximal conductance, $g_{GABA\text{-}B}$, as for the NMDA current, displayed no effect on LAT (**Fig. S5d, left**), but it increased TAU (**Fig. S5d, right**) both in Exc and Inh neurons, because of its slow synaptic action (rise and decay time constants, 90ms and 160ms, respectively) on both neuronal types. Thus, while $g_{GABA\text{-}B}$ possibly accounted for TAU differences in frontal areas and for the absence of change in LAT in Inh neurons, it could not explain the difference in LAT in Exc neurons between LPFC and MCC (**Table S1**).

Interestingly, both NMDA and GABA-B currents, which had no effect at the individual level, were essential at the network scale, suggesting that the influence of slow synaptic transmission on recurrent collective network dynamics are central in determining the time constant TAU in frontal areas.

In summary, one-dimensional network explorations showed that : 1) $g_{AHP}$ was the sole biophysical parameter that changed LAT in Exc but not in Inh, while keeping network activity within the physiological regime ($g_{CAN}$ also changed LAT in Exc, but at the border of network silencing); 2) $g_{GABA-B}$ was the sole biophysical parameter that changed TAU in both Exc and Inh ($g_{CAN}$ and $g_{NMDA}$ also changed TAU, mainly in Exc, but at the border of unrealistic regimes, i.e. network silence and saturation).

Together, these results pointed to $g_{AHP}$ and $g_{GABA-B}$ as major candidates, with the idea that their combined effect in the ($g_{AHP}$, $g_{GABA-B}$) space could account for the differences in temporal signature between the LPFC and the MCC. However, because ionic and synaptic conductances typically display strong non-linear interactions whereby some forms of counter-intuitive compensatory or amplificatory effects can emerge, we nevertheless conducted two-dimensional explorations in the ($g_{AHP}$, $g_{CAN}$), ($g_{AHP}$, $g_{NMDA}$) and ($g_{AHP}$, $g_{GABA-B}$) spaces, with the idea that the relative balance of $g_{AHP}$, which affects LAT, on the one-hand, and of either $g_{CAN}$, $g_{NMDA}$ or $g_{GABA-B}$, which affect TAU, on the other-hand, could synergistically account for both the larger LAT and larger TAU observed in the MCC, compared to the LPFC (**Fig. S6** and **Fig. S3**).

Exploring the ($g_{AHP}$, $g_{CAN}$) space, we found that combined increases of $g_{AHP}$ and $g_{CAN}$ could both 1) increase LAT in Exc neurons but not in Inh neurons (**Fig. S6a**, top) and 2) increase TAU in Exc neurons (**Fig. S6a**, bottom), which translated, quantitatively, as two domains of smaller and larger ($g_{AHP}$, $g_{CAN}$) parameter values that displayed higher similarity to LPFC and MCC data, respectively (**Fig. S6b**; see *Methods*). However, increasing $g_{CAN}$ and $g_{AHP}$ only very weakly varied TAU in Inh neurons (as in one-dimensional explorations), one of the three major changes observed in FS units in the MCC (together with higher LAT and TAU in RS units). While this incapacity marginally reflected in the similarity measure (which integrates similarity in Exc (RS) and FS (Inh) neurons proportional to their relative abundance, i.e. 0.2 for Inh neurons), the model therefore revealed qualitatively insufficient to account for the differences in LPFC and MCC temporal signatures.

The exploration of the ($g_{AHP}$, $g_{NMDA}$) space indicated a situation where combined increases of $g_{AHP}$ and $g_{NMDA}$ could increase LAT in Exc neurons but not in Inh neurons (**Fig. S6c**, top) but could hardly reproduce TAU in Exc and Inh neurons (**Fig. S6c**, bottom), so the qualitative agreement was weak. As a result, quantitatively, the domain of largest MCC similarity displayed modest similarities (**Fig. S6d**). Moreover, as in one-dimensional explorations, $g_{NMDA}$ increased TAU mostly at high values ($g_{NMDA}$ ~ 1), where the network model was near saturation (unrealistic high frequency), while intracellular recordings show no difference in Exc post-synaptic current amplitudes between MCC and LPFC in Monkeys[5].

Contrarily to explorations in ($g_{AHP}$, $g_{CAN}$) and ($g_{AHP}$, $g_{NMDA}$) spaces, exploration in the ($g_{AHP}$, $g_{GABA-B}$) provided two domains of high similarity to LPFC and MCC that indicated a strong quantitative agreement of the model to monkey data (see Main Text). Moreover, these domains were large (relative to the mean values of $g_{AHP}$ and $g_{GABA-B}$ in said domains) indicating robustness to the inherent biological variability present in frontal cortical structures. Finally, qualitative agreement was present, in addition to the quantitative agreement revealed by the similarity measure, in the sense that all three main qualitative differences between LPFC and MCC (with higher LAT and TAU in RS units and a higher TAU in Inh neurons) were well reproduced in this set-up.

| Observable | LAT | | TAU | | Distinct correct effects occur at similar parameter values? | Correct effects occur at realistic network dynamics (silence, saturation)? |
|---|---|---|---|---|---|---|
| Neuron type | Exc. | Inh. | Exc. | Inh. | | |
| Expected LPFC vs MCC difference ? | YES | NO | YES | YES | | |
| Parameter: $g_{CAN}$ | ↓ | NO | ↑ | NO | NO | NO |
| Parameter: $g_{AHP}$ | ↑ | ↓ | NO | NO | YES | YES |
| Parameter: $g_{NMDA}$ | NO | NO | ↑ | ↑ | YES | NO |
| Parameter: $g_{GABA\text{-}B}$ | NO | NO | ↑ | ↑ | YES | YES |

**Table S1. Summary of the effects of the main parameters determining TAU and LAT in the network model.** Expected area differences (LPFC vs MCC) in Exc and Inh neurons are depicted in blue.

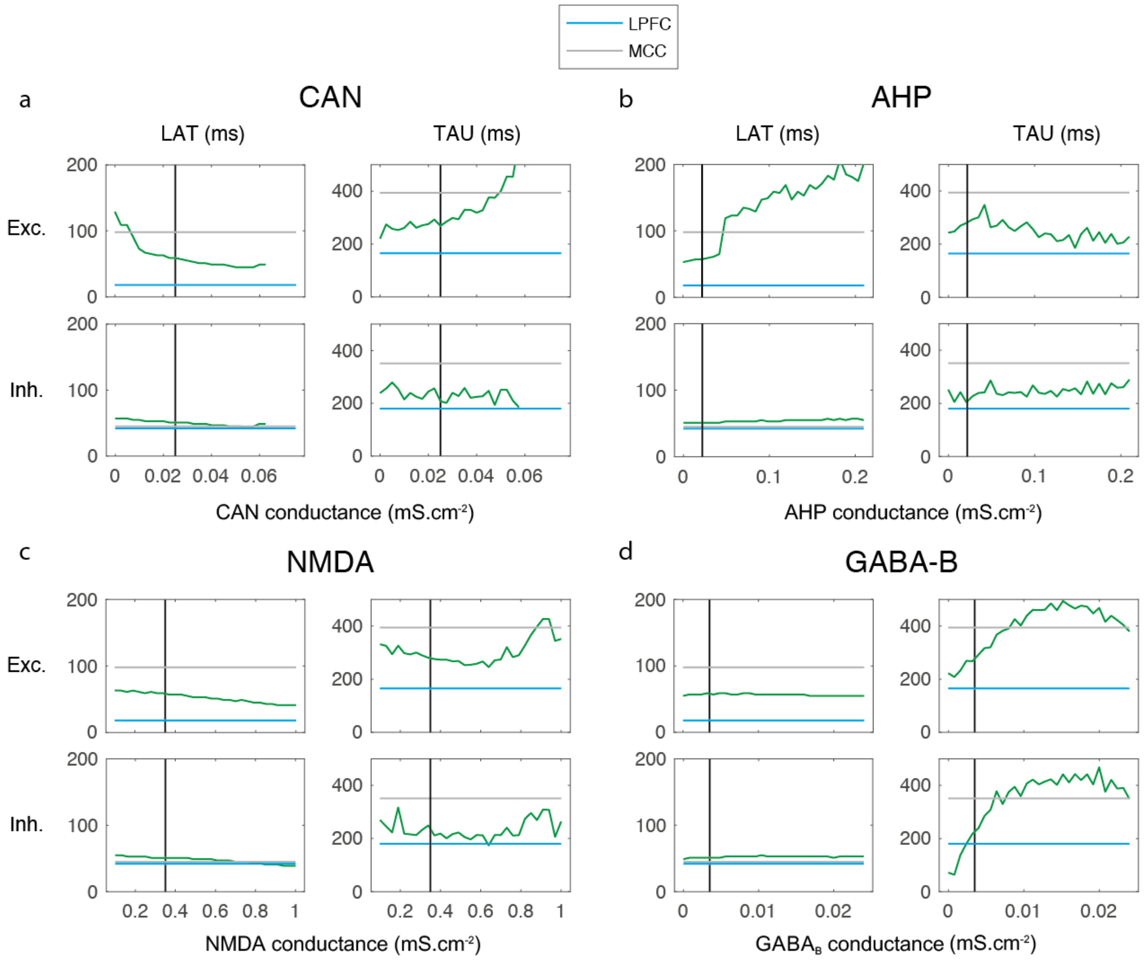


**Figure S5. One-dimensional explorations of key parameters determining TAU and LAT in the network model.** LAT (left) and TAU (right) in Exc (upper) and Inh (lower) neurons in the model (green), as a function of the (**a**) CAN, (**b**) AHP, (**c**) NMDA and (**d**) GABA-B maximal conductances. The default LPFC model parameter value is indicated (black) as well as experimental LAT and TAU in the LPFC (blue) and MCC (grey).

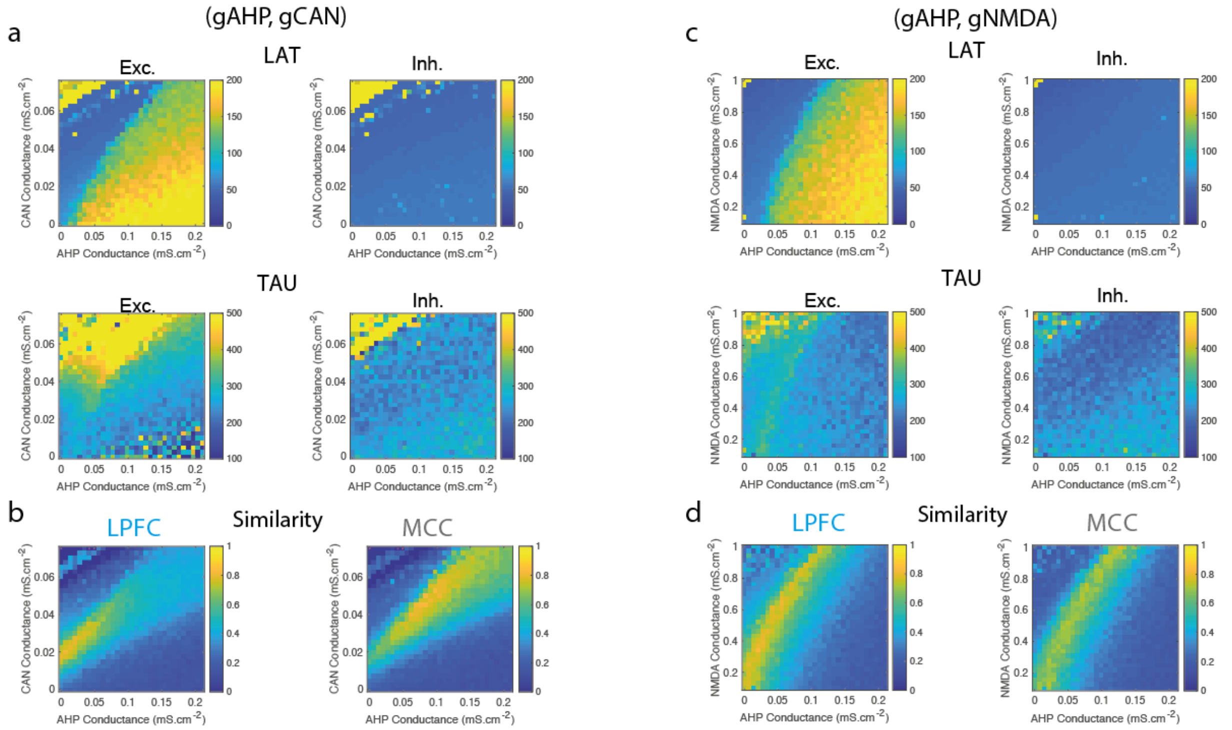


**Figure S6. Two-dimensional explorations in (gAHP, gCAN) and (gAHP, gNMDA) spaces.** (**a**) Mean population LAT (top), TAU (bottom) in Exc (left) and Inh (right) neurons, as a function of AHP and CAN maximal conductances. (**b**) Similarity of the temporal signature between the network model and monkey data in the LPFC (left) and MCC (right), as a function of AHP and CAN maximal conductances (see Methods). (**c**) Same as **(a)** and (**d**) same as **(b)** as a function of AHP and NMDA maximal conductances.

**ISI structure in the network model**

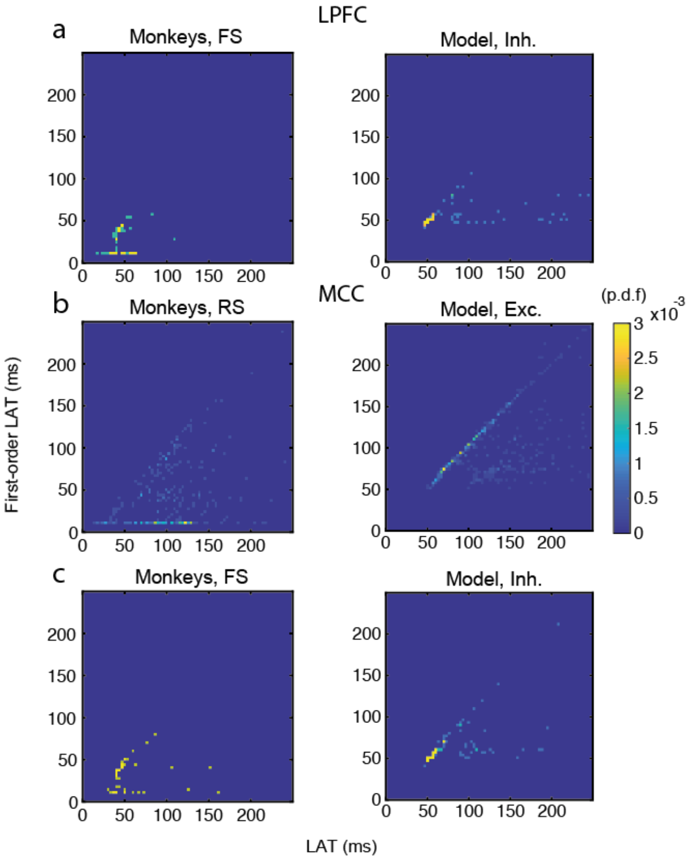


**Figure S7. Relationship between autocorrelogram latency and first-order (ISI) latency in LPFC FS units / inhibitory (Inh) neurons, and in MCC RS units / excitatory (Exc) and MCC FS units / Inh neurons.** Bivariate probability density distribution of the autocorrelogram LAT and first-order latency (i.e. the latency of the ISI distribution) in (**a**) LPFC monkey FS units (left) and network Inh neurons (right), (**b**) MCC monkey RS units (left) and network Exc neurons (right) and (**c**) MCC monkey FS units (left) and network Inh neurons (right).
